# Butyrate rescues chlorpyrifos-induced social deficits through inhibition of class I histone deacetylases

**DOI:** 10.1101/2025.10.19.683261

**Authors:** Leonardo Diaz, Ally Xinyi Kong, Ping Zhang, Jinhua Chi, Khoa Pham, Maja Johnson, Aiden Eno, Isabelle Douglas, Yuxuan Mao, James W. MacDonald, Julia Yue Cui, Theo Bammler, Haiwei Gu, Yijie Geng

## Abstract

Chlorpyrifos (CPF) is a widely used organophosphate pesticide effective through inhibiting acetylcholinesterase, which leads to the accumulation of acetylcholine and continuous nerve stimulation. In addition to its well-known acute toxicity, exposure to CPF has also been linked to chronic conditions such as an increasing risk of autism spectrum disorder (ASD) and adverse effects on gut health, including disturbances to the gut microbiome and metabolism. However, the underlying mechanism of CPF’s contribution to ASD remains unclear, and the roles of the gut microbiome and gut metabolites in CPF-induced neurodevelopmental toxicity remain elusive. Using a high-throughput social behavior assay, we found that embryonic exposure to CPF induced lasting social deficits in zebrafish. Through a small-scale screen of common health beneficial gut microbiome metabolites, we discovered that butyrate effectively rescued CPF-induced social deficits. RNA sequencing of zebrafish brain tissues revealed that early exposure to CPF induced a lasting suppression of neuronal genes, including many ASD risk genes, and elevated expression of circadian genes. Butyrate partially reversed the suppression of key neuronal genes. Butyrate is a non-selective inhibitor of histone deacetylases (HDACs). Through a series of loss-of-function experiments utilizing CRISPR-Cas9-induced knockouts and selective chemical inhibitors, we found that the class I HDAC, HDAC1, most likely mediates butyrate’s rescue effect. Metabolomics analysis detected changes in several nitrogen metabolism-related pathways in the zebrafish gut following CPF exposure. Metagenomics analysis revealed an increase in abundance of the denitrifying bacteria *Pseudomonas* and a reduction in the nitric oxide-sensitive bacteria *Aeromonas* in the CPF-exposed zebrafish gut microbiome. Our results connect CPF-exposure with changes in the gut microbiome, metabolome, epigenetics, gene expression, and behavior, inspiring a novel hypothesis for the underlying molecular mechanisms of CPF-induced neurodevelopmental toxicity. In the long run, our findings may help elucidate how CPF exposure contributes to autism risk and inspire therapeutic developments.

## INTRODUCTION

Chlorpyrifos (CPF) is an organophosphate pesticide critical for agricultural pest management, being used in∼100 countries on 8.5 million acres of crop annually. Its primary mechanism of action involves the inhibition of acetylcholinesterase, which results in acetylcholine accumulation and continuous nerve stimulation, leading to its acute toxicity. Beyond its immediate physiological effects, CPF poses significant environmental risks due to its persistence in soil and water ecosystems, potentially harming non-target species including beneficial insects, birds, aquatic life, and mammals. Recognizing these substantial health and environmental concerns, the Environmental Protection Agency (EPA) has initiated steps to phase out CPF in food production, underscoring the growing scientific consensus about its potential hazards. However, CPF has been continually applied to non-food agriculture, industry, and household uses. In addition to CPF, approximately 40 other organophosphate pesticides remain widely used but are not regulated.

While the acute toxicity of CPF as an acetylcholinesterase (AChE) inhibitor has been well-characterized, the long-term health implications of chronic CPF exposure are not well understood. Chronic CPF exposure has been associated with significant developmental and neurological challenges, particularly in children. Epidemiological studies have documented potential links between CPF exposure and decreased cognitive function, including lower IQ scores and developmental delays^1-3^. Of particular concern is the growing body of evidence suggesting a correlation between CPF exposure and increased risk of autism spectrum disorder (ASD)^4-10^. This potentially contributes to the estimated 40% of autism risk believed to be caused by environmental factors^11,12^.

The potential mechanism underlying this relationship may be linked to CPF’s impact on the gut microbiome.

Exposure to CPF has been shown to induce dysbiosis, or microbial imbalance, in the gastrointestinal tract^13-18^. Pesticide-induced gut dysbiosis is increasingly recognized as a potential contributing factor to neurodevelopmental disorders, including ASD^10,19^. Disruptions in the microbiome can alter metabolite production, which may subsequently influence neurodevelopment through various mechanisms^20,21^. The gut microbiome metabolite butyrate, for example, is a histone deacetylase (HDAC) inhibitor. HDAC inhibitors such as valproic acid can modulate gene expression by preventing the removal of acetyl groups from histones and thereby influencing neurodevelopmental processes. This mechanism is particularly intriguing in the context of ASD, where epigenetic dysregulation is increasingly recognized as a critical factor^22-26^. For example, histone acetylation, a key epigenetic process, has been implicated in ASD pathogenesis^25^. HDAC inhibitors have shown promise in rescuing certain autism-related phenotypes, suggesting a potential therapeutic approach^27^.

Despite these emerging insights, significant knowledge gaps persist. The precise mechanisms by which CPF contributes to ASD remain unclear, and the role of gut microbiome metabolites and epigenetics in CPF-induced neurodevelopmental toxicity remains poorly understood. Our research seeks to address these critical questions through a comprehensive, multi-dimensional approach. Employing a high-throughput social behavior assay in zebrafish, we investigated the effects of embryonic CPF exposure. Through a systematic screening of major gut microbiome metabolites, we discovered that butyrate could effectively rescue CPF-induced social deficits. Mechanistic investigation revealed that the classes I HDAC, HDAC1, likely mediates butyrate’s rescue effect. We further probed the molecular mechanisms underlying CPF’s neurodevelopmental toxicity using multi-omics analyses, including metagenomics, metabolomics, and RNA sequencing. RNA sequencing results revealed significant downregulation of neuronal genes, many of which have been previously associated with ASD. Butyrate partially reversed this downregulation. Metabolomics and metagenomics analysis results both revealed changes related to nitrogen metabolism and nitric oxide (NO) production. These results inspire a hypothesis that CPF induces social deficit by first enhancing NO production through alterations in gut microbiome composition, followed by NO-induced selective inhibition of class I HDACs, which ultimately causes imbalanced histone acetylation and dysregulated neuronal gene expression.

## RESULTS

### Butyrate rescued chlorpyrifos-induced social deficit in zebrafish

Social behavioral deficit is a defining characteristic of ASD. Due to CPF’s association with ASD, we examined whether CPF can induce social deficits in zebrafish using a high-throughput behavioral platform, Fishbook^28^. We found that embryonic exposure to CPF at 0-3 days post fertilization (dpf) induced social deficits at the juvenile stage (25 dpf) in a dose dependent manner (Figure 1A & Supplementary Figure S1A). 15 µM CPF consistently induced significant social deficit and was used for all subsequent rescue experiments. Exposure to CPF is known to cause gut microbiome dysbiosis and alter the production of beneficial gut microbiome metabolites^13,17,18,29^. To test the hypothesis that CPF-induced social deficit may be alleviated by beneficial gut microbiome or liver metabolites, we conducted a small-scale screen of 7 metabolites, including butyrate (in the form of sodium butyrate), acetic acid, chenodeoxycholic acid (CDCA), cholic acid, 3-indolepropionic acid (IPA), succinic acid, and ursodeoxycholic acid (UDCA), all applied at a 10 µM concentration (Figure 1B). Only butyrate successfully rescued CPF-induced social deficit (Figure 1B). To confirm this finding, we applied sodium butyrate at concentrations ranging from 1.25 µM to 40 µM at 2-fold increments and observed a bell-shaped rescue curve which peaked at 10 µM, validating that 10 µM butyrate can most effectively rescue CPF-induced social deficit (Figure 1C).

**Figure 1.**
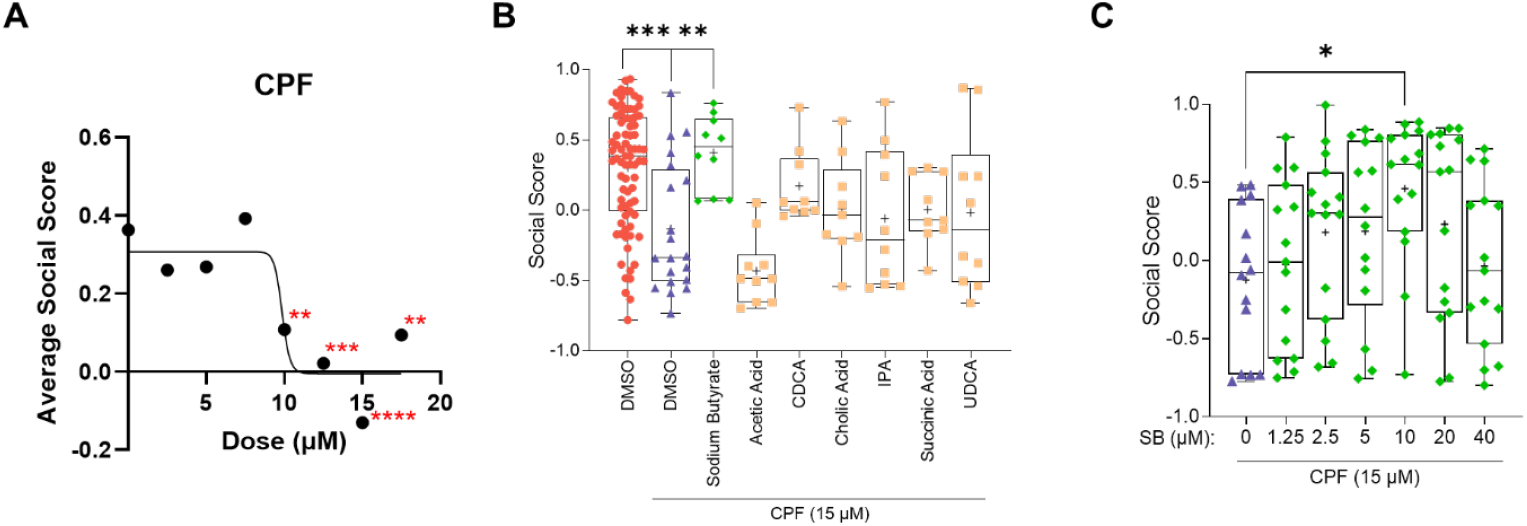
Butyrate rescued CPF-induced social deficit in zebrafish. (**A**) Chlorpyrifos (CPF) induced social deficits through embryonic exposure in a dose-dependent manner. CPF’s effects were significant from 10-17.5 µM and peaked at 15 µM. (**B**) Selected gut microbial metabolites and liver metabolites were tested at 10 µM for their abilities to rescue social deficits. Colored dots represent the social scores of individual fish. Sodium butyrate effectively rescued social deficits induced by 15 µM CPF. UDCA: ursodeoxycholic acid; IPA: indole-3-propionic acid; CDCA: chenodeoxycholic acid. (**C**) Sodium butyrate (SB) rescued social deficits induced by 15 µM CPF in a dose-dependent manner. Significant rescue was observed at 10 µM. Significance was calculated by one-way ANOVA and Dunnett’s multiple comparison test. \**p*<0.05, \*\**p*<0.01, \*\*\**p*<0.001, \*\*\*\**p*<0.0001.

### Inhibition of class I histone deacetylases phenocopied butyrate’s rescue effect

Histone deacetylases (HDACs) are a family of enzymes that can remove acetyl groups from lysine residues on histone proteins, thereby altering chromatin structure and gene expression. HDACs are classified into five major classes based on sequence homology and functional properties, including classes I, IIa, IIb, III (sirtuins), and IV. Butyrate is a pan-inhibitor of HDACs. To identify the HDACs involved in butyrate’s rescue effect, we tested several class-specific HDAC inhibitors including valproic acid (VPA; a class I & IIa Inhibitor), trichostatin A (TSA; a class I, II, & IV Inhibitor), and nicotinamide (NAM; a class III / sirtuins inhibitor). Compounds were applied to juvenile zebrafish overnight prior to Fishbook assay; test subjects were pre-exposed to CPF at 0-3 dpf. We found that only the class I & IIa Inhibitor VPA rescued CPF-induced social deficit (Figure 2A).

**Figure 2.**
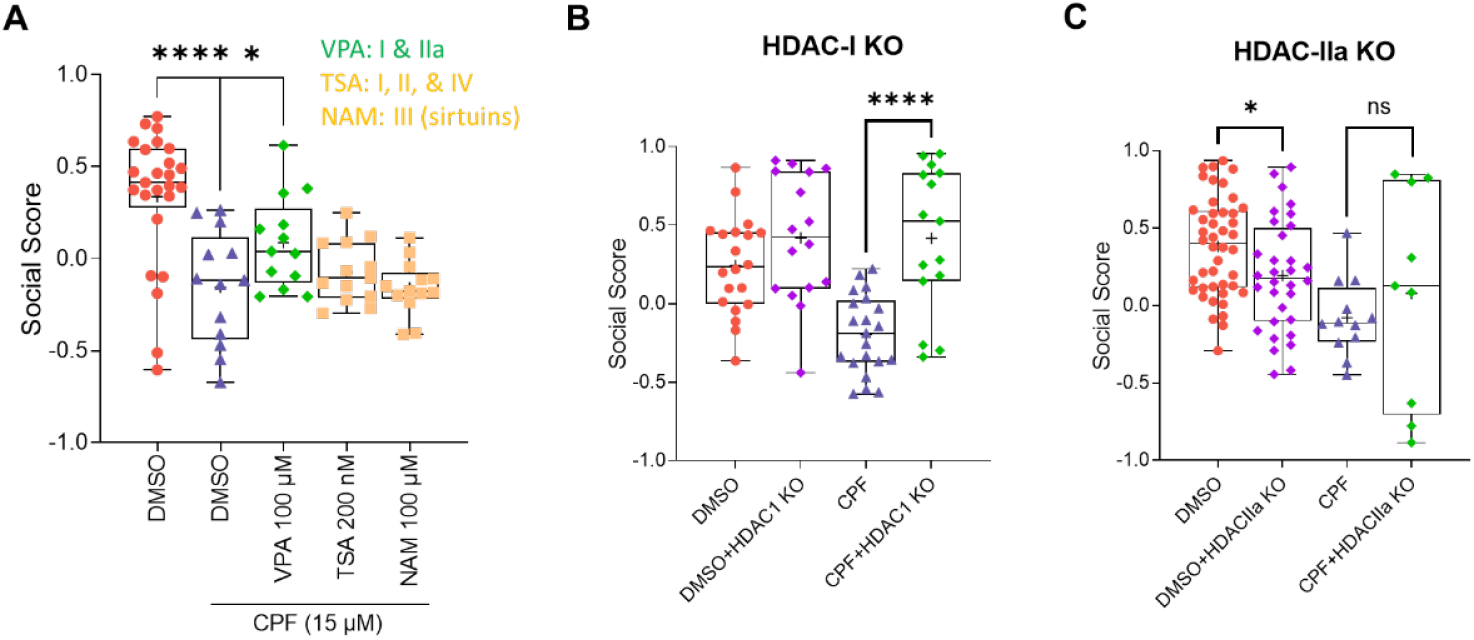
Class I HDAC inhibition phenocopied butyrate’s rescue effect. (**A**) Valproic acid (VPA) (100 µM) but not trichostatin A (TSA) (200 nM) or nicotinamide (NAM) (100 µM) rescued social deficits induced by 15 µM CPF. VPA is an inhibitor of classes I and IIa HDACs, TSA is an inhibitor of classes I, II, and IV HDACs, and NAM is an inhibitor of class III (sirtuins) HDACs. Significance was calculated by one-way ANOVA and Dunnett’s multiple comparison test. (**B**) CRISPR-Cas9 induced F0 knockout (KO) of zebrafish class I HDACs (HDAC-I) rescued social deficits induced by 15 µM CPF. This experiment shares the same DMSO and CPF controls with the HDAC1 KO experiment shown in Figure 3A. Significance was calculated by two-way ANOVA with Fisher’s LSD test for post hoc analysis. (**C**) CRISPR-Cas9 induced F0 KO of zebrafish class IIa HDACs (HDAC-IIa) failed to rescue social deficits induced by 15 µM CPF. Significance was calculated by two-way ANOVA with Fisher’s LSD test for post hoc analysis. ns: not significant, \**p*<0.05, \*\*\*\**p*<0.0001.

To further zoom in on the biological pathway underlying butyrate and VPA’s rescue effects, we attempted to simultaneously knock down all class I HDACs or all class IIa HDACs in zebrafish, respectively, and evaluate their impacts on sociality with or without pre-exposure to CPF. In humans, class I HDACs include HDAC1, HDAC2, HDAC3, and HDAC8. The zebrafish genome possesses orthologs genes for the human *HDAC1 (hdac1), HDAC3 (hdac3)*, and *HDAC8 (hadc8)*, but not *HDAC2. HDAC1, HDAC3, and HDAC8* are all expressed in the brain. The human class IIa HDACs include HDAC4, HDAC5, HDAC7, and HDAC9. HDAC7 is not expressed in the brain and is therefore excluded from subsequent research. The zebrafish genome also possesses orthologs genes for the human *HDAC4* (*hdac4*), *HDAC5* (*hdac5*), and *HDAC9* (*hdac9*). To knockout all genes in each class using CRISPR-Cas9, we pooled 3 guide RNAs for each gene in the same class (*HDAC1/3/8* for class I, and *HDAC4/5/9* for class IIa), and separately co-injected each guide RNA mixture with Cas9 protein into 1-2 cell stage zebrafish embryos to induce knockout. We found that F0 knockout of class I HDACs phenocopied butyrate and VPA by robustly rescuing CPF-induced social deficit (Figure 2B), whereas the class IIa knockout did not rescue CPF-induced social deficit and instead inhibited social behavior when compared to wild-type DMSO control (Figure 2C). Together, these results suggest that class I HDACs are involved in the regulation of CPF-induced social deficit in zebrafish.

### Inhibition of HDAC1 alone effectively rescued chlorpyrifos-induced social deficit

To identify the HDAC enzyme specifically involved in social behavioral regulation following CPF pre-exposure, we used CRISPR-Cas9 to individually knock out the zebrafish orthologs of each class I HDAC gene. We found that knocking out *hdac1* robustly upregulated social behavior in vehicle control fish as well as CPF-exposed fish (Figure 3A). Knocking out *hdac3* or *hdac8* both modestly rescued social behavior in CPF-exposed fish, but did not boost sociality in the control fish (Figures 3B & 3C). In fact, *hdac8* knockout reduced social score when compared to wild-type DMSO control (Figure 3C). We also attempted to rescue CPF-induced social deficit via overnight exposure to selective HDAC inhibitors that specifically target HDAC1 (BRD-6929), HDAC3 (RGFP966), and HDAC8 (PCI-34051). We found that the HDAC1 selective inhibitor, BRD-6929, strongly rescued CPF-induced social deficit in a dose-dependent manner, with 50 µM BRD-6929 most robustly boosted social behavior (Figure 3D & Supplementary Figure S1B). The HDAC3 selective inhibitor RGFP966 modestly improved social behavior in CPF-exposed fish, especially when applied at 50 µM, but did not consistently boost the average social score to above 0 at higher dosages tested (Figure 3E & Supplementary Figure S1C): a social score below 0 indicates social avoidance, a social score of 0 indicates no social preference, and a social score above 0 indicates social interest^28^. The HDAC8 selective inhibitor PCI-34051 failed to rescue CPF-induced social deficit at the dosages tested (Figure 3F & Supplementary Figure S1D). These results suggest that HDAC1, and to a lesser extent, HDAC3, likely mediate butyrate and VPA’s rescue effects, and that targeted inhibition of HDAC1 alone is sufficient to robustly rescue social deficit induced by CPF exposure.

**Figure 3.**
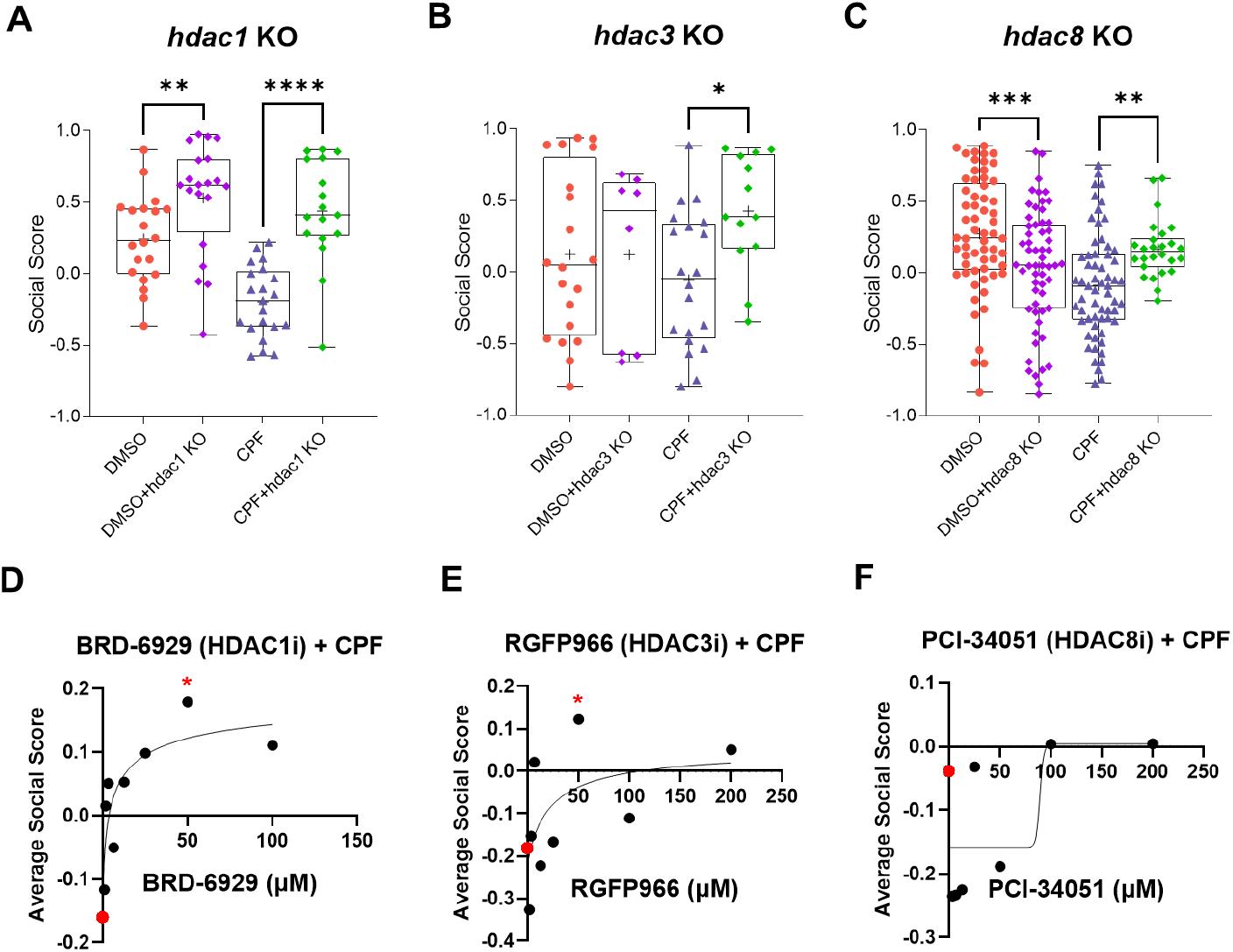
Selective inhibition of HDAC1 effectively rescued CPF-induced social deficit. (**A**) CRISPR-Cas9 induced F0 KO of the zebrafish *hdac1* gene robustly rescued social deficits induced by 15 µM CPF (CPF+*hdac1* KO). Social behavior in DMSO control was also significantly boosted by *hdac1* KO (DMSO+*hdac1* KO). This experiment shares the same DMSO and CPF controls with the HDAC-I KO experiment shown in Figure 2B. Significance was calculated by two-way ANOVA with Fisher’s LSD test for post hoc analysis. (**B**) Knocking out the zebrafish *hdac3* gene modestly rescued social deficits induced by 15 µM CPF (CPF+*hdac3* KO). Significance was calculated by two-way ANOVA with Fisher’s LSD test for post hoc analysis. (**C**) Knocking out the zebrafish *hdac8* gene rescued social deficits induced by 15 µM CPF (CPF+*hdac8* KO). Significance was calculated by two-way ANOVA with Fisher’s LSD test for post hoc analysis. (**D-F**) Dose dependent rescue of CPF-induced (15 µM CPF) social deficits following overnight exposures of BRD-6929, a selective inhibitor of HDAC1 (HDAC1i) (D), RGFP966, a selective inhibitor of HDAC3 (HDAC3i) (E), and PCI-34051, a selective inhibitor of HDAC8 (HDAC8i) (F). Red dot marks the DMSO control of each experiment. The HDAC1 inhibitor BRD-6929 robustly rescued CPF-induced social deficit in a dose-responsive manner (D). The HDAC3 inhibitor RGFP966 modestly rescued social deficit in a dose-responsive manner (E). The HDAC8 inhibitor PCI-34051 failed to rescue CPF-induced social deficit. Significance was calculated by one-way ANOVA and Dunnett’s multiple comparison test. \**p*<0.05, \*\**p*<0.01, \*\*\**p*<0.001, \*\*\*\**p*<0.0001.

### Butyrate partially rescued the selective inhibition of neuronal genes by chlorpyrifos

To explore the molecular mechanisms underlying CPF-induced social deficits and butyrate’s rescue effect, we conducted whole-brain RNA-sequencing (RNA-seq) using 25 dpf zebrafish across four experimental treatments, including DMSO control treatment (DMSO), sodium butyrate treatment (SB), CPF treatment (CPF), and sodium butyrate rescue of CPF-exposed fish (CPF+SB). CPF and DMSO were exposed to embryonic and early-larval-stage zebrafish at 0-3 dpf. Butyrate was applied the day before sample collection and kept overnight. Zebrafish gene names were converted to the names of their human orthologs for pathway analysis. Gene Set Enrichment Analysis (GSEA) using Gene Ontology’s (GO) Biological Process (BP) terms found that embryonic exposure to CPF significantly inhibits expression of neuronal genes related to neuronal projection (axonal and dendritic), synaptogenesis (presynaptic and postsynaptic), and learning, as compared to the DMSO control (Figure 4A & Supplementary Figures S2-S4). Out of the top 20 significantly downregulated GO pathways, 13 are related to the regulation of neuronal processes (Figure 4A). Genes associated with butanoate (butyrate) metabolism are negatively enriched in CPF-treated zebrafish compared to the control fish (Supplementary Figure S5), although this pathway did not make the cut for the top 20 downregulated pathways ranked by normalized enrichment scores (NESs). Significantly downregulated neuronal genes belonging to the 13 pathways form a compact protein-protein interaction (PPI) network (Supplementary Figure S6). Within this network, the top 30 hub genes were identified by network topology analysis and ranked by node degree values (the number of direct interactions within the network). These hub genes again form a compact PPI network (Figure 4B). While CPF suppressed the expression of these neuronal hub genes, butyrate restored their expressions in CPF-exposed fish, as shown by a gene expression heatmap (Figure 4C) and statistical analysis of the average normalized expression levels of these genes (Figure 4D). Approximately half (14 out of 30) of the hub genes are high-confidence or suspected ASD risk genes as identified by the Simons Foundation Autism Research Initiative (SFARI) in their list of SFARI genes^30,31^, including *BRAF*^32^ (SFARI gene score [SGS]: 1S), *CAMK2B*^33^ (SGS: S), *CDH2*^34^ (SGS: 3S), *DRD2*^35^ (SGS: 2), *GRIA2*^36^ (SGS: 1), *GRIA3*^37^ (SGS: S), *GRIN1*^38^ (SGS: 1), *GRIN2B*^39^ (SGS: 1), *GRM5*^40^ (SGS: 2), *MAPK3*^41^ (SGS: 2), *NLGN1*^42^ (SGS: 2), *PAX6*^43^ (SGS: S), *SNAP25*^44^ (SGS: 2), and *SYP*^45^ (SGS: 3). The genes *GRIA2* (adjusted *p* value [*padj*] = 0.00098), *GRIA3* (*padj* = 0.0011), and *GRIN1* (*padj* = 0.013) were significantly upregulated by butyrate (Figure 4C: asterisks). These results indicate that a network of neuronal genes associated with ASD risk are selectively downregulated in CPF-exposed zebrafish, and this downregulation is partially rescued by butyrate treatment.

**Figure 4.**
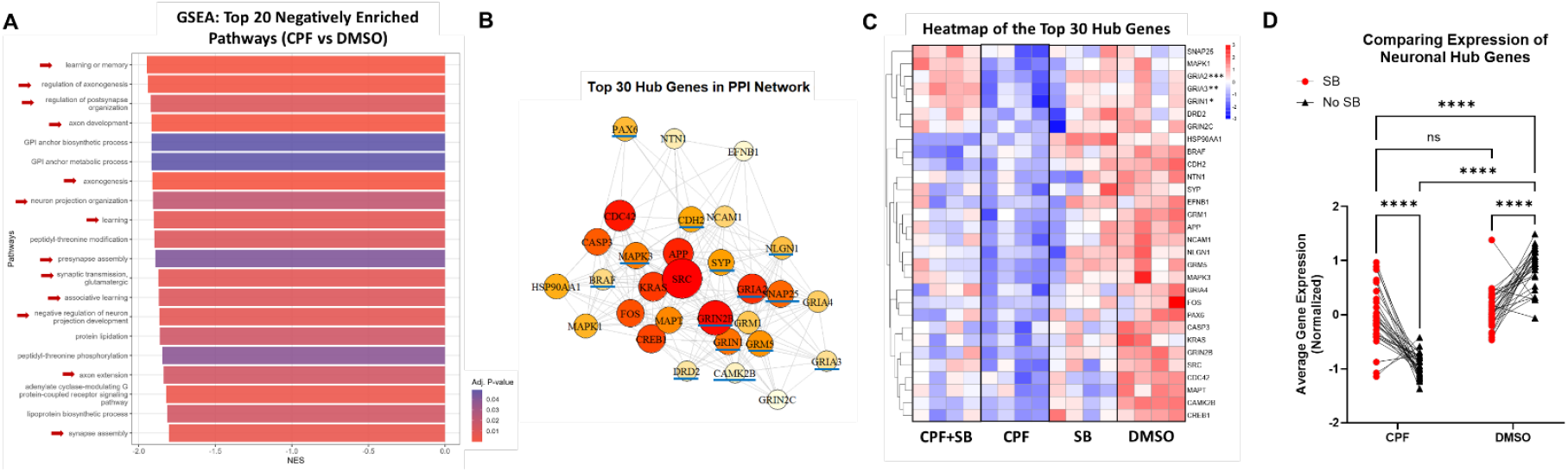
Butyrate partially rescued sustained downregulation of neuronal genes induced by early exposure to CPF. (**A**) Gene Set Enrichment Analysis (GSEA) of RNA-seq data using Gene Ontology’s (GO) Biological Process (BP) terms identified 13 pathways (red arrows) related to neuronal projection, synaptogenesis, and learning among the top 20 downregulated pathways in CPF-treated samples as compared to the DMSO control samples. Significantly downregulated pathways (*p* < 0.05) are ranked by Normalized Enrichment Score (NES). (**B**) Hub genes were identified by network topology analysis of a protein-protein interaction network formed by protein products of significantly downregulated neuronal genes found in the CPF samples compared to the control samples. All genes belong to the 13 neuronal pathways shown in (A). The top 30 hub genes were ranked by node degree values. Colors represent hub gene rankings, with red denoting the top-ranking genes. The size of each colored circle represents the degree centrality (number of connections in the network). Blue underlines mark the SFARI ASD risk genes. (**C**) Gene expression heatmap shows butyrate (SB) partially reversing the downregulation of 30 neuronal hub genes induced by CPF-exposure (CPF+SB vs. CPF). (**D**) Comparing the average normalized expressions of 30 neuronal hub genes. Neuronal hub genes are significantly downregulated in CPF:No-SB samples compared to the DMSO:No-SB controls. Butyrate (SB) significantly upregulated the expression of neuronal hub genes in CPF-treated fish (CPF:SB), although not enough to match the level of expression in DMSO:No-SB control samples. Gene expression levels are not significantly different in CPF:SB samples as compared to DMSO:SB samples. Lines between treatment groups connect expression levels of the same gene. Significance was calculated by two-way ANOVA with Tukey’s multiple comparison test. ns: not significant, \**p*<0.05, \*\**p*<0.01,\*\*\**p*<0.001, \*\*\*\**p*<0.0001.

### Embryonic exposure to chlorpyrifos induced sustained overexpression of key circadian genes

GSEA using GO-BP terms found that 5 out of the 20 positively enriched pathways are associated with circadian regulation (Figure 5A & Supplementary Figure S7). RNA-seq data demonstrated a significant overexpression of circadian genes in the juvenile zebrafish brain: 11 out of the top 21 significantly upregulated genes are circadian genes, including *per1a, per1b, per2, cry1a, cry1b, cry2, cry5, nr1d1, ciarta, bhlhe41*, and *si:ch211-132b12*.*7* (Figure 5B). Because circadian gene expressions oscillate during the day, to prevent RNA-seq sample collection times affecting gene expression levels and bias the result, we repeated the same experimental treatments and collected brain samples from CPF-treated fish and control fish at the exact same time of the day through brain dissections performed in parallel by two experienced experimenters. Quantitative PCR analysis using these samples confirmed circadian gene upregulation in CPF-treated fish compared to the control fish (Figure 5C). It is worth noting that the human orthologs of several identified circadian genes are known ASD risk genes in the SFARI gene list, including *PER1*^46^ (SGS: 2), *PER2*^47^ (SGS: 2), and *NR1D1*^48^ (SGS: 2).

**Figure 5.**
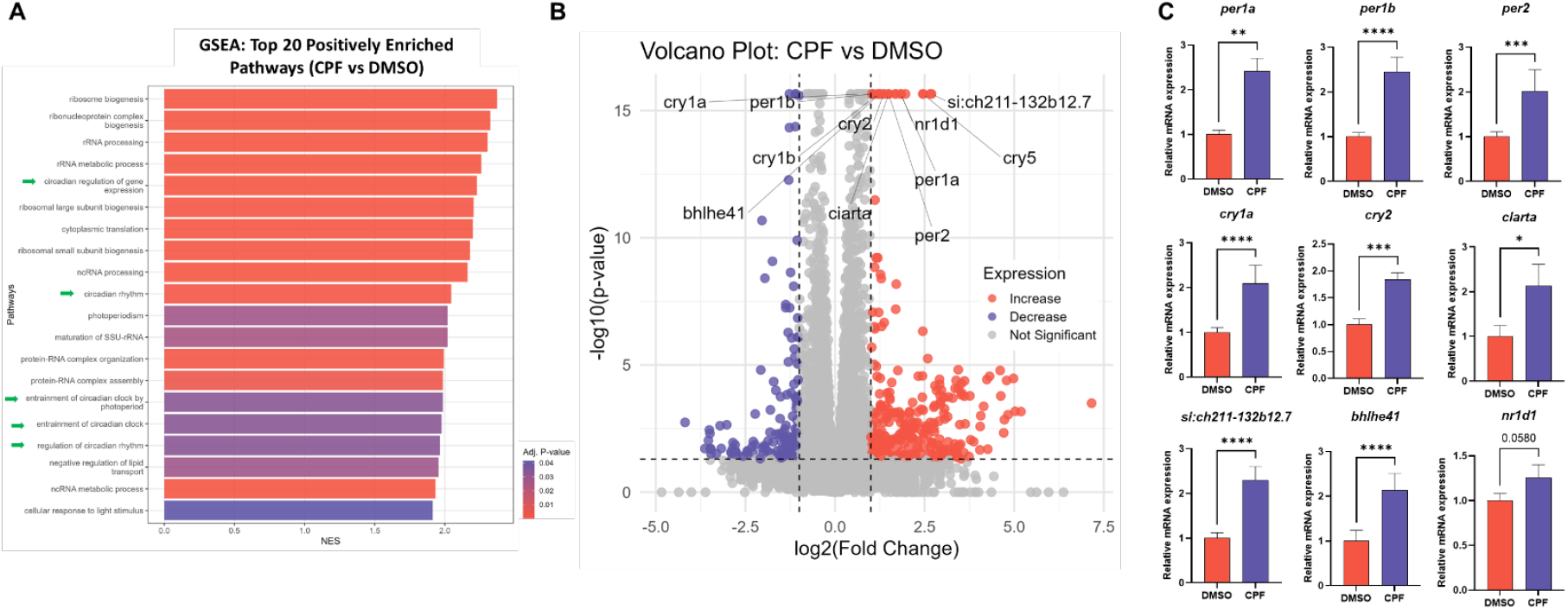
Early CPF exposure induced sustained overexpression of key circadian genes. (**A**) Gene Set Enrichment Analysis (GSEA) of RNA-seq data using Gene Ontology’s (GO) Biological Process (BP) terms. Showing the top 20 upregulated pathways in CPF-treated samples as compared to the DMSO control samples, among which 5 are related to circadian regulation (green arrows). Significantly upregulated pathways (*p* < 0.05) are ranked by Normalized Enrichment Score (NES). (**B**) Volcano plot showing the RNA-seq result comparing gene expression in CPF-treated samples and DMSO control samples. 11 of the top 21 significantly upregulated genes (ranked by adjusted *p* value) are circadian genes. (**C**) Quantitative PCR analyses conducted using samples from an independent experimental replicate validated RNA-seq results by confirming upregulation of key circadian genes following embryonic exposure to CPF. CPF-treated fish and DMSO control fish were dissected at the exact same time during the day by two experimenters working side-by-side. Significance was calculated by two-tailed Student’s *t* test. \**p*< 0.05, \*\**p*<0.01, \*\*\**p*<0.001, \*\*\*\**p*<0.0001.

Significantly upregulated circadian genes belonging to the five identified circadian pathways (Figure 5A) form a compact PPI network (Supplementary Figures S8A & S8B). Plotting an expression heatmap of these circadian hub genes using RNA-seq data showed that butyrate treatment following CPF-exposure (CPF+SB) further boosted the expression of these key circadian genes compared to the CPF-treated sample (Supplementary Figure S8C). Statistical analysis of the average normalized gene expression levels of circadian hub genes confirmed this elevation in gene expression in the CPF+SB samples compared to the CPF samples (Supplementary Figure S8D). This observation was also verified by analyzing the differential gene expression data between CPF+SB and CPF samples using GSEA, which found that 6 out of the 10 upregulated pathways are associated with circadian regulation (Supplementary Figure S8E). These results demonstrate that butyrate did not reverse CPF-induced upregulation of circadian gene expression, but instead further boosted their expression. We thus reason that butyrate’s social behavioral rescue effect is unlikely mediated by the circadian pathway.

### Chlorpyrifos induced lasting changes in nitrogen metabolism pathways in the gut

To test the hypothesis that CPF-exposure may alter the production of beneficial metabolites by the gut microbiome, we again exposed zebrafish to three treatment conditions: DMSO, CPF, and CPF+SB. CPF and DMSO were exposed to zebrafish at 0-3 dpf. Butyrate was applied the day before sample collection and kept overnight. The entire gastrointestinal tract, including the gut contents, were collected from juvenile stage zebrafish (25 dpf) through dissection. Untargeted metabolomics were conducted to determine the metabolite profiles of each sample. Enrichment analysis using MetaboAnalyst 6.0^49^ against the “Feces” dataset in the Human Metabolome Database^50,51^ (HMDB) detected similar patterns of metabolite profile changes in CPF-treated samples compared to fecal samples collected from individuals with inflammatory bowel disease (IBD), including Crohn’s disease and colitis, irritable bowel syndrome (IBS), and autism (Supplementary Figure S9A). We also ran several enrichment analyses against the Relational Database of Metabolomics Pathways^52,53^ (RaMP-DB), the Small Molecule Pathway Database^54^ (SMPDB), and the Kyoto Encyclopedia of Genes and Genomes^55^ (KEGG) to detect metabolomic pathways altered in the CPF samples as compared to the DMSO controls. We consistently identified urea cycle, arginine metabolism, and aspartate metabolism among the top enriched pathways (Figures 6A-6C). Both arginine and aspartate are intermediate metabolites of the urea cycle. Interestingly, changes in the nitric oxide (NO) pathway and nitric oxide synthase (NOS) activity were detected by surveying RaMP-DB (Figure 6A). L-arginine is a precursor for the endogenous production of NO through NOS, thus connecting urea cycle, arginine metabolism, and aspartate metabolism with the NO/NOS pathway (Figure 6D). Enrichment analysis using SMPDB detected significant changes in urea cycle, arginine metabolism, and aspartate metabolism between the CPF+SB and CPF samples (Figure 6E), indicating that these pathways were also altered by butyrate treatment. However, none of the three pathways passed significance threshold when surveying the KEGG and RaMP-DB databases for metabolic changes between the CPF+SB and CPF samples (Supplementary Figures S9B & S9C), suggesting that the butyrate-induced changes in these pathways are likely mild.

**Figure 6.**
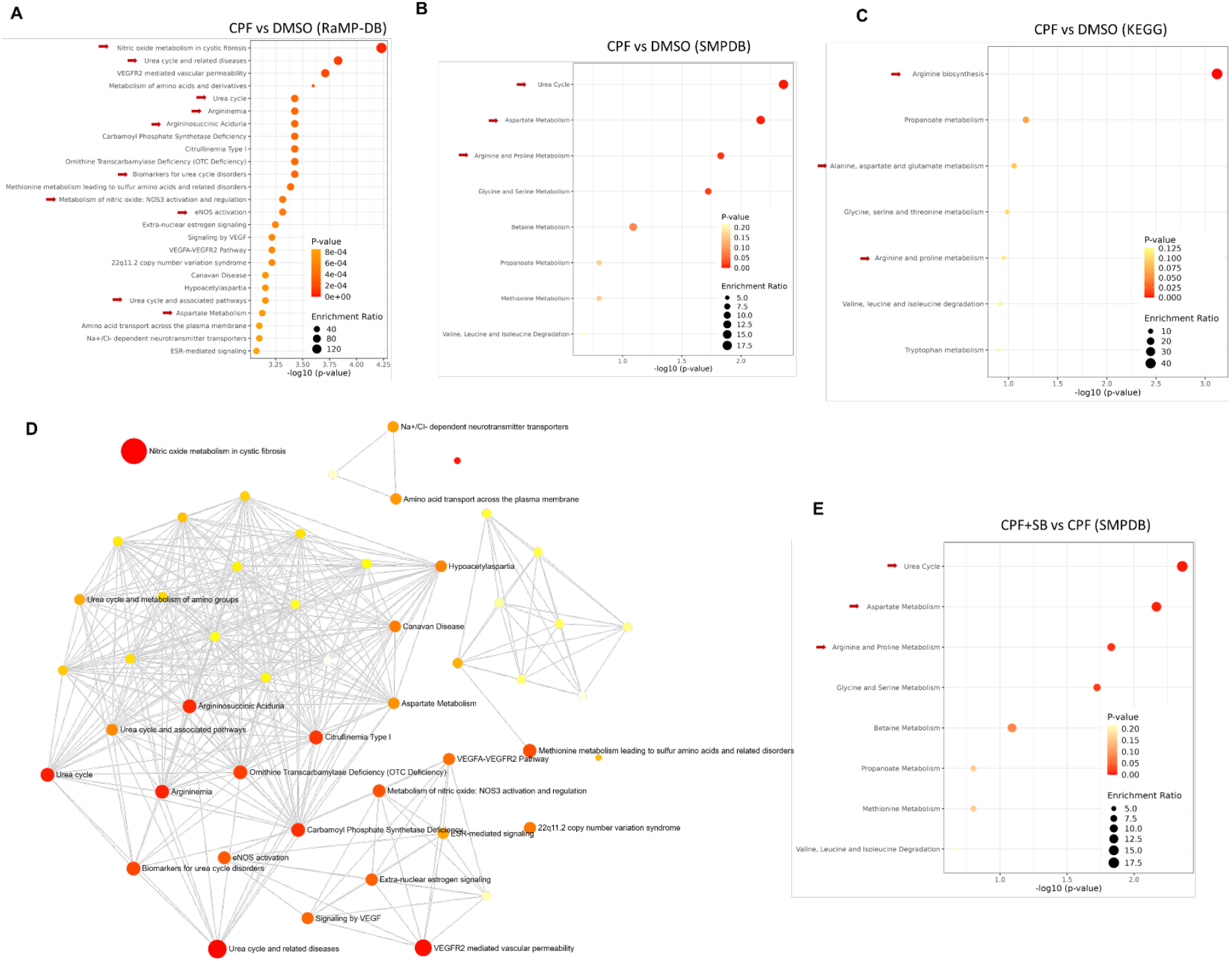
Early CPF exposure selectively impacted nitrogen metabolism-related pathways in the juvenile zebrafish gut. (**A-C**) Enrichment analyses detected changes in gut metabolomic pathways in CPF-treated fish compared to DMSO control fish, based on untargeted metabolomics analysis of gastrointestinal tissues and gut contents. Surveying the RaMP-DB (A), SMPDB (B), and KEGG (C) databases consistently identified urea cycle, arginine metabolism, and aspartate metabolism among the top enriched pathways (red arrows). Both arginine and aspartate are intermediate metabolites of the urea cycle. Changes in nitric oxide (NO) metabolism and nitric oxide synthase (NOS) activity were detected by surveying RaMP-DB (A). Color coding represents levels of significance (*p* value). The size of each dot represents enrichment ratio. (**D**) Network view of the enrichment analysis result in (A). L-arginine is a precursor for the endogenous production of NO through NOS, thereby connecting arginine metabolism, aspartate metabolism, and urea cycle with the NO/NOS pathway. (**E**) Enrichment analysis comparing metabolomic changes between CPF+SB and CPF samples based on untargeted metabolomics analysis of gastrointestinal tissues and gut contents. Pathways related to the urea cycle, arginine metabolism, and aspartate metabolism (red arrows) were identified by surveying SMPDB.

### Chlorpyrifos increased the abundance of denitrifying bacteria in the zebrafish gut microbiome

We then conducted metagenomics analysis to examine if changes in the zebrafish gut microbiome contributes to its metabolomic alterations following CPF exposure. CPF and DMSO were exposed to zebrafish at 0-3 dpf. Following ∼ 3 weeks of growth in nursery, fecal matters were collected from juvenile stage zebrafish (25 dpf) for DNA extraction and shotgun metagenomic sequencing. Sankey diagrams were generated to show combined taxonomy of species found in the DMSO control and CPF samples, respectively (Figures 7A & 7B). We conducted differential analyses for abundance (i.e., number of observations for a given taxonomic level) and prevalence (i.e., presence/absence of bacteria at a given taxonomic level) between the two treatment groups at every taxonomic level, from species to kingdom. We only identified differentially abundant taxa and found no evidence for differentially prevalent taxa. The count of significant taxa at each level is shown in Figure 7C. There are two differentially abundant phyla, *Actinomycetota* and *Cyanobacteriota* (Supplementary Figure S10), each increased in abundance by 2-to 2.5-fold following CPF exposure, as shown by their β coefficient values (Supplementary Figure S10B). At the class level, *Actinomycetes* and *Cyanophyceae* were found to have similar 2-to 2.5-fold increases in abundance in the CPF samples (Supplementary Figure S11). The order *Micrococcales* and the family *Microbacteriaceae* both increased by ∼2.5-fold in abundance following CPF exposure (Supplementary Figures S12 & S13). Although no significant changes in abundance were detected at the genus level, 139 species were found to be differentially abundant (Figure 7C & Supplementary Table S1), among which 86 species belong to the genus *Microbacterium* (Figure 7D & Supplementary Figure S14A). All differentially abundant *Microbacterium* species showed an increase in abundance in the CPF-treated samples as compared to the DMSO control. A boxplot comparing normalized abundance measures for the significant and non-significant *Microbacterium* species found that the non-significant species have similar yet much lower abundance in CPF and DMSO samples, which may explain why even though so many *Microbacterium* species are differentially abundant, when combined at the genus level the difference is no longer significant (Supplementary Figure S14B). The *Microbacterium* genus is classified under the family *Microbacteriaceae*, the order *Micrococcales*, the class *Actinomycetes*, and the phylum *Actinomycetota*.

**Figure 7.**
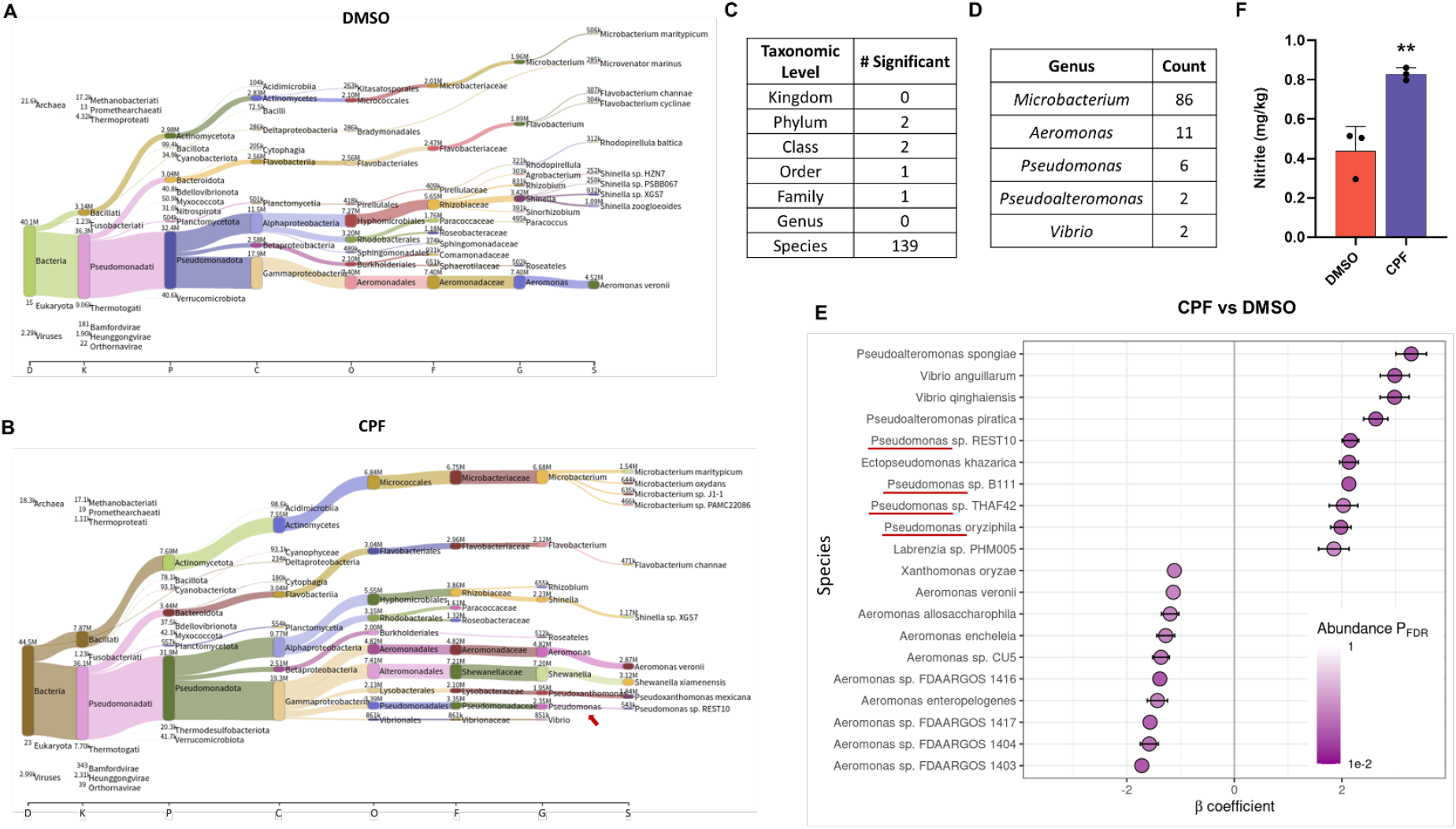
Early CPF exposure led to enrichment of denitrifying bacteria in the juvenile zebrafish gut microbiome. (**A**) Gut microbial taxonomic structure of the DMSO samples. Kraken generated read counts were classified at each level of the taxonomic tree and summed as we go up the taxonomy. (**B**) Gut microbial taxonomic structure of the CPF samples. Red arrow points to the genus *Pseudomonas* which is increased in abundance in the CPF samples as compared to the DMSO control samples. (**C**) Number of taxa that are differentially abundant at each taxonomic level. (**D**) Selected counts of differentially abundant species by genus. (**E**) β coefficient estimates and significance for the 10 species with the largest beta values (log fold change) and the 10 species with the smallest beta values. β coefficients are calculated for the test between CPF and DMSO samples. Error bars represent the confidence interval of each coefficient. Color denotes false discovery rate (FDR). There are a total of 139 significant species, only the 20 with the most extreme beta values are shown in this plot. (**F**) Nitrite concentration was significantly elevated in 6 dpf zebrafish larvae following embryonic (0-3 dpf) treatment of CPF. n=85 larvae per condition. Significance was calculated by two-tailed Student’s *t* test. \*\**p*<0.01.

In addition to the 86 *Microbacterium* species, 11 *Aeromonas* species, 6 *Pseudomonas* species, 2 *Pseudoalteromonas* species, and 2 *Vibrio* species were found to be differentially abundant in CPF and DMSO samples (Figure 7D). When plotting species with the 10 highest and 10 lowest β coefficient values, no *Microbacterium* species were found (Figure 7E). Instead, we found the 2 *Pseudoalteromonas* species, 2 *Vibrio* species, and 4 out of the 6 *Pseudomonas* species among the top 10 species with the highest increase in abundance (Figure 7E). In the Sankey diagram for CPF samples (Figure 7B), the genus *Pseudomonas* shows the highest read count (2.35M) (Figure 7B: red arrow) compared to *Pseudoalteromonas* (165k; not shown by the plot) and *Vibrio* (851k), indicating that *Pseudomonas* is a dominant genus among the top species with increased abundance following CPF exposure. Many *Pseudomonas* species are facultative anaerobes that carry out complete denitrification – the stepwise reduction of nitrate (NO_3_^−^) to nitrite (NO_2_^−^), nitric oxide (NO), nitrous oxide (N2O), and finally nitrogen gas (N2). Studies of activated sludge and biofilm reactors show that among denitrifying bacteria, *Pseudomonas* is frequently the dominant genus^56,57^. This unique metabolic capability connects changes in the abundance of *Pseudomonas* with the observed changes in nitrogen metabolism in the CPF samples (Figures 6A-6D). Some members of the genera *Pseudoalteromonas* and *Vibrio* have also been reported to be capable of at least partial denitrification^58-60^. *Microbacterium* is generally not known to possess denitrifying activity. In contrast to the abovementioned genera, all 11 *Aeromonas* species decreased in abundance following CPF exposure (Supplementary Table S1). Of the top 10 species showing the greatest reduction in abundance in the CPF samples compared to controls, 9 are *Aeromonas* (Figure 7E). This may be due to *Aeromonas’* susceptibility to the antimicrobial effect of nitric oxide (NO)^61-64^, a metabolite presumably produced at a higher amount by the increased denitrifying gut bacteria in CPF-treated fish. Indeed, using Griess assay, we detected significantly elevated nitrite levels in 6 dpf zebrafish larvae pre-exposed to CPF from 0-3 dpf (Figure 7F). As a stable oxidation product of NO, nitrite levels are commonly used as an indirect measurement of NO. Elevated nitrite levels therefore indicate elevated NO levels in zebrafish following CPF exposure. This result is consistent with the identification of disrupted NO/NOS-related pathways through metabolomics analysis (Figures 6A & 6D). The delay between CPF exposure (0-3 dpf) and the detection of elevated NO (6 dpf) suggest a lasting effect of CPF on NO production, which is likely at least in part mediated by changes in gut microbiome composition.

### A NO-HDAC hypothesis on CPF-induced social deficits

These findings inspired us to propose a working hypothesis that CPF induces ASD-relevant neurodevelopmental defects through excessive NO production and the resulting imbalanced activities of class I HDACs. We hereafter refer to it as the NO-HDAC hypothesis of CPF’s neurodevelopmental toxicity (Figure 8A). Briefly, we propose that CPF exposure first increases NO production through a combination of three (likely independent) mechanisms: (1) our results show that via an unknown mechanism, CPF exposure increases the abundance of denitrifying bacteria in the gut microbiome, most notably *Pseudomonas* (Figure 7E), which could enhance NO production (Figure 7F) through the denitrification pathway; (2) CPF exposure has been found to trigger the expression of inducible nitric oxide synthase (iNOS) in various experimental systems, resulting in increased NO production^65-69^; and (3) CPF prevents the breakdown of the neurotransmitter acetylcholine (ACh) via its activity as an irreversible inhibitor of the enzyme acetylcholinesterase (AChE), which can in turn activate endothelial nitric oxide synthase (eNOS) and thereby promoting NO production^70-72^. NO is known to inhibit HDAC2 (not present in zebrafish) and HDAC8 through S-nitrosylation of two cysteine residues which are conserved in all class I HDACs^73-75^, yet through an unknown mechanism, HDAC1 and HDAC3 are largely immune to S-nitrosylation despite possessing the same conserved cysteine residues^76^. We therefore reason that an abnormally high level of NO induced by CPF exposure will lead to excessive (compared to physiological level) inhibition of HDAC8 through S-nitrosylation, while HDAC1 and HDAC3 activities will remain relatively intact due to their resistance to S-nitrosylation. This selective inhibition of HDAC2/8 over HDAC1/3 can skew the normal physiological balance of class I HDAC activities in the brain, altering histone acetylation patterns in a way that suppresses the expression of neuronal genes critical for social behavioral development and thus leading to the social deficit phenotype.

**Figure 8.**
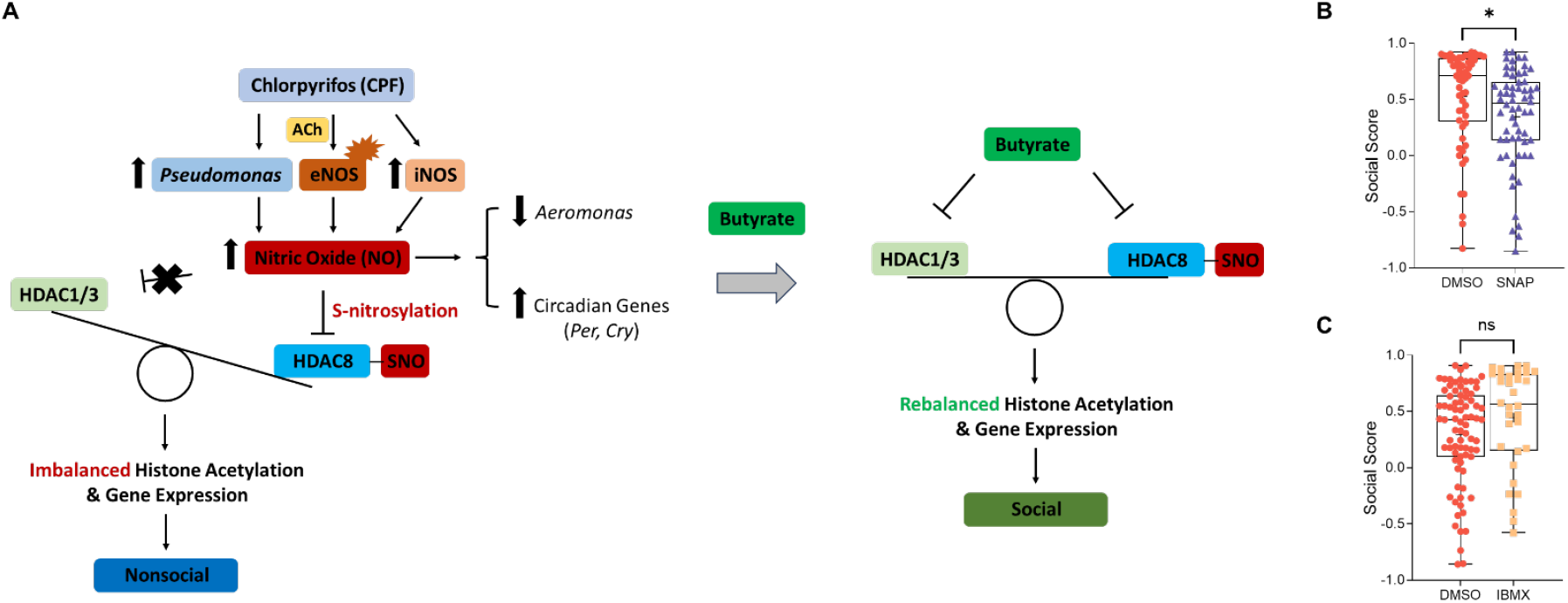
A working hypothesis for CPF’s neurodevelopmental toxicity. (**A**) A schematic demonstration of the NO-HDAC hypothesis. Chlorpyrifos (CPF) exposure increases the abundance of the denitrifying bacteria *Pseudomonas* in the gut microbiome, enhancing the production of nitric oxide (NO) in the gut through the denitrification pathway. CPF is also known to stimulate NO production by activating eNOS through acetylcholine (Ach) accumulation and promoting iNOS expression. Elevated NO levels result in the selective inhibition of HDAC8 through S-nitrosylation. HDAC1 and HDAC3 are resistant to S-nitrosylation and thus escape inhibition by NO. This selective inhibition skews the balance of class I HDAC activity in the developing zebrafish brain, altering histone acetylation patterns in a way that suppresses the expression of neuronal genes critical for social behavior, thereby leading to social behavioral deficits. Other biological effects of NO production include the downregulation of NO-sensitive bacteria of the genus *Aeromonas* and the upregulation of circadian genes, particularly the *Per* and *Cry* family genes. Butyrate inhibits all members of class I HDACs, including the zebrafish HDAC1, HDAC3, and HDAC8, thereby resetting the genome-wide balance of histone acetylation at gene loci targeted by class I HDACs. SNO: S-nitrosothiols. (**B**) Embryonic exposure (0-3 dpf) to the NO donor SNAP (10 µM) induced social deficits in Fishbook assay. Significance was calculated by two-tailed Student’s *t* test. \**p*< 0.05. (**C**) Embryonic exposure (0-3 dpf) to the non-selective PDE inhibitor IBMX (100 µM) did not induce measurable social deficits in Fishbook assay. Significance was calculated by two-tailed Student’s *t* test. ns: not significant.

This model helps to explain butyrate’s rescue effect: based on the NO-HDAC hypothesis, we propose that butyrate rescues CPF-induced social deficit by simultaneously inhibiting all members of class I HDACs, including the zebrafish hdac1, hdac3, and hdac8, thus correcting the genome-wide imbalance of histone acetylation by “resetting” and rebalancing the global histone acetylation states, especially at gene loci targeted by class I HDACs. This rebalancing theory also explains why knocking out or pharmaceutically inhibiting the zebrafish hdac1 and hdac3 resulted in successful rescue of the CPF-induced social deficit phenotype (Figures 3A, 3B, 3D, & 3E), whereas chemical inhibition of hdac8 failed to rescue the deficit (Figure 3F). Knocking out *hdac8* did rescue social deficit induced by CPF (Figure 3C), but we reason that this is likely due to the loss of S-nitrosylation target for NO. Knocking out *hdac8* significantly inhibited social behavior compared to wild-type control (Figure 3C), while in contrast, *hdac1* knockout robustly boosted social behavior (Figure 3A). In comparison, *hdac3* knockout did not significantly affect social behavior when compared to wild-type control (Figure 3B), indicating that *hdac1* and *hdac8* may be the two members of class I HDACs that are key to maintaining the hypothesized state of histone acetylation balance. Finally, the elevated NO in our NO-HDAC model also explains the reduction of the NO-sensitive bacteria genus *Aeromonas* (Figure 7E) and the upregulation of circadian genes (Figure 5), particularly the *Per* and *Cry* family genes^77-79^, following CPF exposure. A more in-depth analysis of the NO-HDAC hypothesis can be found in the Discussion section.

As a preliminary test of this hypothesis, we exposed zebrafish embryos to the NO donor S-nitroso-N-acetylpenicillamine (SNAP). Fishbook assay detected a lasting deficit in social behavior at the juvenile stage following embryonic exposure to SNAP (Figure 8B), suggesting a role of NO in social behavioral development, which is in line with our model’s prediction. A major biological effect of NO is to promote the production of cyclic guanosine monophosphate (cGMP), an effect that can be pharmacologically mimicked by inhibiting cGMP-degrading phosphodiesterases (PDEs). Embryonic exposure to 3-isobutyl-1-methylxanthine (IBMX), a broad-spectrum PDE inhibitor, did not induce social deficits in zebrafish (Figure 8C), indicating that NO likely modulates social behavior through a cGMP-independent mechanism, which again aligns with our model (Figure 8A).

## DISCUSSION

Our study reveals a multi-layered mechanism through which embryonic exposure to CPF, a commonly used organophosphate pesticide, induces lasting social behavioral deficits in zebrafish, a core phenotype relevant to ASD. Using high-throughput behavioral screening, transcriptomics, metabolomics, and metagenomics, we uncovered that CPF disrupts social behavior by altering the gut microbiome and gut-derived metabolites, leading to suppression of neuronal gene expression in the brain. Notably, we found that butyrate, a gut microbiome metabolite and HDAC inhibitor, rescues CPF-induced social deficits and inhibition of neuronal genes. Mechanistic dissection identified HDAC1 as a key mediator of this rescue effect. Additionally, we observed sustained overexpression of circadian genes and significant alterations in nitrogen-related metabolic pathways, including upregulation of denitrifying bacteria such as *Pseudomonas* and depletion of nitric oxide (NO)-sensitive *Aeromonas* species.

The NO-HDAC hypothesis (Figure 8A) developed based on these findings is strongly supported by our data and helps to explain observations in our study. First, CPF-treated zebrafish show significant increase in *Pseudomonas* abundance, an established genus of denitrifying bacteria capable of producing NO under anaerobic and microaerobic conditions^80-86^. Second, concurrent with the increase in *Pseudomonas* species, we observed significant enrichment in nitrogen metabolism-related pathways, including arginine, aspartate, and urea cycle metabolism, which are tightly linked to NO synthesis and may represent secondary effects of increasing denitrification and NO production mediated by *Pseudomonas*. Third, CPF exposure caused a reduction in *Aeromonas* species which are facultative anaerobes known to be sensitive to NO^61-64^, suggesting elevated NO levels in the gut environment. Fourth, brain transcriptome analysis revealed a pronounced downregulation of neuronal genes, many of which are high-confidence ASD risk genes. Butyrate rescued both the CPF-induced downregulation of neuronal genes and social deficits. More selectively, HDAC1 inhibition was sufficient to robustly rescue behavioral phenotypes, indicating that HDAC1 is a pivotal component of this pathway. Finally, elevated expression of circadian genes observed in our study, in particular *Per* and *Cry* family genes, is consistent with the known ability of NO to induce these genes via CREB phosphorylation and S-nitrosylation of Bmal1^77-79^. Butyrate failed to reverse the CPF-induced circadian gene upregulation, suggesting that circadian gene expression changes are likely not causal for behavioral deficits but are rather a secondary effect of NO signaling.

Our hypothesis is also supported by extensive literature. CPF has been shown to elevate NO levels in the brain *in vivo*, across multiple animal models. Rodent and fish studies consistently demonstrate increased NO metabolites and iNOS activity following CPF exposure^66,68,87^. Furthermore, other organophosphates such as malathion^88^, parathion^89^, diazinon^90^, and dichlorvos^87^ exhibit similar effects, pointing to a conserved mechanism of NO elevation induced by this class of chemicals. NO inhibits HDAC2 and HDAC8 through the S-nitrosylation of two cysteine residues that are highly conserved among all members of class I HDAC^73-75^. However, HDAC1 and HDAC3 are not subjected to NO-mediated S-nitrosylation despite possessing the same conserved cysteine residues^76^. HDAC2 is not present in the zebrafish genome, leaving HDAC8 to be the sole target of NO among members of class I HDACs. Class I HDACs play essential regulatory roles in the development of the central nerves system^91^. In particular, HDAC8 is a SFARI gene (SGS: S) found to be associated with Cornelia de Lange syndrome^92-94^, autism^34,95-97^, Rett-related disorder^98^, and intellectual disability^99,100^. Our hypothesis proposes that NO-mediated S-nitrosylation of HDAC8 leads to increased histone acetylation and gene activation at specific gene loci, disrupting the balanced levels of histone acetylation and gene expression among key neuronal genes, which is typically maintained by all class I HDACs in a coordinated manner. Butyrate and VPA presumably rescued this imbalanced state by simultaneously inhibiting all class I HDACs, thereby resetting histone acetylation for all class I HDAC targeted regions of the genome. This theoretical framework also explains the rescue effects of inhibiting HDAC1 and HDAC3 (Figures 3A-3B & 3D-3E). The rescue effect of knocking out *hdac8* (Figure 3C) may be a result of NO losing its target molecule and therefore no longer able to modulate gene expression through HDAC8 S-nitrosylation. Our approach to rescue neurodevelopmental deficits through epigenetic reprogramming echoes our previous success in correcting social deficits induced by imbalanced histone methylation using a similar strategy^28^. The hypothesis of differential inhibition of class I HDACs by NO also resonates well with the concept of a post-translational code for class I HDACs^101^.

Importantly, elevated nitric oxide levels and nitrosative stress have been associated with the pathogenesis of ASD^102-104^. Studies found increased concentrations of nitric oxide, its stable metabolites (e.g., nitrite and nitrate), and biomarkers of nitrosative stress in the plasma and saliva of children diagnosed with ASD^105-108^. The same phenomenon was observed in animal models of ASD including *Shank3*-deficient mice^109-111^ and *Cntnap2* mutant mice^111^. *Shank3*-deficient mice exhibited widespread changes in the S-nitroso-proteome, in particular the S-nitrosylation of key synaptic proteins^109^, proteins involved in glutamate transmission^110^, and protein products of high-risk SFARI genes^110^. Inhibition of NO production by a neuronal nitric oxide synthase (nNOS) inhibitor restored the reduced synaptic protein expression and decreased dendritic spine density in *Shank3* mutant mice^111^. In *Cntnap2* mutant mice, nNOS inhibition similarly reversed the increase in NO metabolites and nitrosative stress markers, and rescued deficits in synaptic protein expression and dendritic spine density^111^. These findings suggest that dysregulated nitric oxide signaling may be a common feature in autism, providing additional support for our model that links CPF-induced NO elevation to neurodevelopmental and behavioral alterations.

Additional research is warranted to further test our hypothesis. First, direct quantification of NO and its metabolites (e.g., nitrite/nitrate) in the brain and gut of CPF-exposed zebrafish would validate the proposed increase in nitrosative stress. Second, experimental detection of S-nitrosylation in the zebrafish HDAC1, HDAC3, and HDAC8 enzymes following CPF-exposure would verify the hypothesize differential regulations of class I HDACs by NO. Third, genome-wide profiling of histone acetylation, particularly at neuronal gene loci, could confirm CPF-induced epigenetic dysregulation and butyrate-mediated rescue. Fourth, pharmacological or genetic inhibition of NO production (e.g., using NOS inhibitors) should be tested for their ability to reverse social deficits in CPF-exposed animals. If successful, this would establish NO as a functional mediator between microbiome alterations and neurodevelopmental outcomes.

In conclusion, our findings bridge CPF-induced microbiome dysbiosis, NO production, epigenetic modification, and ASD-relevant behavior through a novel mechanistic axis involving denitrifying bacteria and selective HDAC regulation. This integrative framework not only sheds light on CPF’s contribution to neurodevelopmental risk but also provides a compelling rationale for exploring NO signaling and HDAC1 as therapeutic targets for environmentally triggered neurodevelopmental disorders.

## MATERIALS AND METHODS

### Zebrafish husbandry

Zebrafish were housed at 26°C–27°C on a 14-hour light, 10-hour dark light cycle. Wild-type AB strain was used for all experiments. All zebrafish experiments were approved by the Institutional Animal Care and Use Committee at the University of Washington.

### Chemical exposure

Fertilized embryos were sorted and transferred into 100 mm diameter and 15 mm deep petri dishes at 100 embryos per dish. Each dish was filled with 25 mL of HEPES-buffered E3 medium. Compound stocks were prepared in DMSO and stored at -20°C. Vehicle controls receiving an equal volume of DMSO as the corresponding compound treatments. For embryonic compound treatment, embryos were exposed to compounds from 0-3 dpf. Dead embryos were removed at 1 and 2 dpf to prevent contamination. At 3 dpf, all viable larvae were rinsed with E3 medium and transferred into clean petri dishes containing fresh E3 medium. At 5 dpf, larvae from each petri dish were transferred to separate nursery tanks and raised to 25 dpf for Fishbook assay. For overnight compound treatment, 24 dpf fish were transferred into 100 mm diameter and 15 mm deep petri dishes at 15 fish per dish and exposed to chemicals overnight (∼15 hours). Following the overnight exposure, fish were rinsed and tested in the Fishbook assay.

The following compounds were also obtained from Cayman Chemical (Ann Arbor, MI, USA): DMSO (CAS # 67-71-0, Item No. 20131), chlorpyrifos (CAS # 2921-88-2, Item No. 21412), sodium butyrate (CAS #156-54-7, Item No. 13121), 3-indolepropionic acid (IPA, CAS #830-96-6, Item No. 28821), cholic acid (CAS #81-25-4, Item No. 20250), chenodeoxycholic acid (CDCA, CAS #2646-38-0, Item No. 35346), ursodeoxycholic acid (UDCA, CAS #31687-65-7, Item No. 15121), valproic acid (VPA, CAS #1069-66-5, Item No. 13033), nicotinamide (CAS #98-92-0, Item No. 11127), trichostatin A (TSA, CAS #58880-19-6, Item No. 89730), SNAP (CAS # 67776-06-1, Item No. 82250), and IBMX (CAS # 28822-58-4, Item No. 13347). Acetic acid (CAS #64-19-7, Product No. 45754) and succinic acid (CAS #110-15-6, Product No. S9512) were obtained from Sigma-Aldrich. The following HDAC inhibitors were obtained from MedChemExpress (Monmouth Junction, NJ, USA): BRD-6929 (CAS #849234-64-6, Cat. No.: HY-100719), RGFP966 (CAS #1357389-11-7, Cat. No.: HY-13909), and PCI-34051 (CAS #950762-95-5, Cat. No.: HY-15224).

### Fishbook assay

The Fishbook assay was run as described previously^28^. Briefly, the test arena consists of a total of 44 3D printed, 10-mm-deep, 8.5-mm-wide, and 80-mm-long rectangular chambers grouped together in parallel with each other. Each chamber is divided into three compartments by two transparent acrylic windows (1.5 mm thick): a 60-mm-long middle testing chamber to place the test subject and two 8.5-mm-long end chambers to place the social stimulus fish or remain empty, respectively. Test subjects were each placed inside an individual test chamber using a plastic transfer pipette with its tip cut off to widen the opening. A 3D printed white comb-like structure was placed in front of the social stimulus compartment to block the test subject’s visual access to social stimulus fish before testing begins. After test subjects were placed inside the chambers, the arena was placed inside the imaging station, and the combs were removed to visually expose the social stimulus fish to the test subjects. Following a brief acclimation period, a 10-min test session was video recorded.

Videos were streamed through the software Bonsai^112^. Videos were analyzed in real time during recording, and the frame-by-frame x and y coordinates of each fish relative to its own test compartment were exported as a CSV file. Data was analyzed using custom Python scripts to calculate social scores and generate tracking plots. Social score was defined as a fish’s average y-axis position for all frames. The middle of each test chamber was designated as the origin of the y axis, with an assigned value of zero. A value of 1 was assigned to the end of the chamber next to the social stimulus fish and a value of −1 to the other end of the chamber next to the empty control compartment. In this coordinate system, all social scores have values between −1 and 1. A higher social score demonstrates a shorter average distance between a test subject and a social stimulus fish during a test, which suggests a stronger social preference.

Fishbook results were analyzed and plotted using GraphPad Prism. For analysis of multiple groups, normal distribution of datasets was examined using Shapiro-Wilk test to confirm that more than half the data was normally distributed and standard deviations fell within the variance ratio. To compare multiple groups with one independent variable, one-way analysis of variance (ANOVA) assuming gaussian distribution and equal standard deviation was performed, using Dunnett’s test to correct for multiple comparisons. To compare groups with two independent variables, two-way ANOVA was performed followed by Fisher’s Least Significant Difference (LSD) test for post hoc analysis. *P* values less than 0.05 were considered significant.

### CRISPR-Cas9 gene knockout

We designed 3 sets of sgRNAs targeting each HDAC ortholog to maximize the efficiency of F0 knockout^113-115^. CHOPCHOP^116^ was used for the sgRNA design. sgRNAs targeting early coding exons were selected to introduce nonsense mutations as early as possible and increase the likelihood of loss-of-function mutations. All gRNA sequences are shown in Supplementary Table S2.

The SP6 promoter sequence was added before the target RNA sequence, followed by an overlap adapter, which is complementary to the 5’ end of an 80 bp constant oligo according to a published protocol^117^. Oligonucleotides were synthesized by Eurofins Scientific. Double-stranded DNAs were generated by Phusion Hot-Start Flex DNA Polymerase (New England Biolabs) using the gene specific oligos and the constant oligo. The double-stranded DNA was cleaned up using the Zymo Clean and Concentrator-5 kit (Zymo Research). *In vitro* sgRNA transcription was conducted using the MEGAscript SP6 transcription kit (Thermo Fisher Scientific) and cleaned up using the Zymo RNA Clean and Concentrator-5 kit (Zymo Research).

To perform zebrafish embryonic microinjections, pairs of male and female adult zebrafish were kept in a mating cage overnight while separated by a divider. 1-cell-stage embryos were collected the next day early in the morning immediately before injection by pulling out the divider to allow breeding. Injection solution was prepared by combining sgRNA with 2 µM Spy Cas9 NLS and 1X NEBuffer (New England Biolabs).

### Zebrafish brain and intestine dissection

At 25 days post-fertilization, zebrafish were euthanized by immersion in icy water for 15 minutes in petri dishes. For brain dissection, the fish were positioned vertically, dorsal side up, in a pre-carved slot within dissection wax on a petri dish. The lower half of the body was immobilized using Gorilla Glue to ensure stability. The petri dish was filled with ice-cold phosphate-buffered saline (PBS) to prevent desiccation during the procedure. Brains were carefully extracted from the skull using the tip of a sharp syringe needle. Five brains were collected per replica, with four replicas prepared per condition. For intestine dissection, the fish were positioned horizontally on a clean petri dish, with the head and tail secured using glue. Ice-cold PBS was added to maintain hydration, and intestines were gently isolated using a dissection scalpel. Twenty-four intestines were collected per replica, with four replicas prepared per condition. Immediately after harvest, both brains and intestines were flash-frozen in liquid nitrogen and stored at -80°C.

### RNA-sequencing

Total RNA was isolated from frozen 25 dpf zebrafish brains using Quick-RNA Miniprep Kit (Zymo Research, Irvine, California) by following the manufacturer’s protocol. RNA concentrations were quantified using a NanoDrop 1000 Spectrophotometer (Thermo Scientific, Waltham, Massachusetts) at 260 nm. Sample QC was performed with Agilent 4150 bioanalyzer to identify samples with RIN>7. 200ng qualified RNA from each sample was processed for library preparation. In brief, mRNA enrichment was performed on total RNA using oligo(dT)-attached magnetic beads. The enriched mRNA with poly(A) tails was fragmented using a fragmentation buffer, followed by reverse transcription using random N6 primers to synthesize cDNA double strands. The synthesized double stranded DNA was then end-repaired and 5’-phosphorylated, with a protruding ‘A’ at the 3’ end forming a blunt end, followed by ligation of a bubble-shaped adapter with a protruding ‘T’ at the 3’ end. The ligation products were PCR amplified using specific primers. The PCR products were denatured to single strands, and then single-stranded circular DNA libraries were generated using a bridged primer. The constructed libraries were quality-checked and sequenced after passing the quality control. The library was amplified with phi29 to make DNA nanoball (DNB) which had more than 300 copies of one molecular. The DNBs were load into the patterned nanoarray and pair end 150 bases reads were generated in the way of sequenced by synthesis. The sequencing was conducted on the MGI T7 platform.

Raw sequencing reads in FASTQ format were first processed to quality control using FastQC (v0.11.9), followed by Clumpify (BBMap, v38.90), Trimmomatic (v0.39) and fastp (v0.23.2) on Ubuntu 20.04 LTS to obtain clean reads. Analysis of the clean reads were performed using R Studio (v4.4.1). Clean reads were first aligned to the reference genome (*Danio rerio*, GCF_000002035.6_GRCz11_genomic). A gene expression matrix was created using featureCounts from the Rsubread (v2.18.0) package in R Studio. Differential expression analysis was performed using the Limma R package (v3.60.6). Genes with *p*-value < 0.05 and |log2FoldChange| ≥ 1 were considered significantly differentially expressed (DEGs). Volcano plot and heatmap were generated using EnhancedVolcano (v1.22.0), pheatmap (v1.0.12) and ggplot2 (v3.5.1) packages.

### Gene set enrichment analysis (GSEA)

DEGs were subjected to functional enrichment analysis using the clusterProfiler R package (v4.12.6). GSEA was performed using the gseGO() and gseKEGG() functions for GO and KEGG enrichment respectively. Gene IDs were converted using biomaRt (v2.60.1) to ensure compatibility across databases. Pathways with adjusted p-value < 0.05 and normalized enrichment score (NES) > 1 were considered significant.

### Protein–Protein Interaction (PPI) Analysis

To explore protein interaction relationships among DEGs, PPI networks were constructed using the STRING database (v12.0, https://string-db.org). Then PPI interaction data were imported into R studio for further network analysis. Network topology analysis was performed using igraph (v2.3.0) package to identify hub genes. Node degree (the number of direct interactions) was calculated for each gene, and genes with the highest degree values were considered hub genes.

### Quantitative PCR

Total RNA samples extracted from zebrafish brains were reverse-transcribed into cDNA using SuperScript™ IV First-Strand Synthesis System per the manufacturer’s protocol. (ThermoFisher Scientific, Waltham, Massachusetts). The resulting cDNA products were amplified by qPCR, using SYBR Green qPCR Master Mix (GlpBio Technology Inc, Montclair, California) in a Bio-Rad CFX Connect Real-Time System (Bio-Rad, Hercules, California). Data was normalized to the housekeeping gene Beta-actin. Primers were designed using the National Center for Biotechnology Information (NCBI) Primer-BLAST and synthesized by Eurofins Genomics (Louisville, Kentucky). Primer sequences are shown in Supplementary Table S3.

### Untargeted metabolomics by LC-MS

Acetonitrile (ACN), methanol (MeOH), ammonium acetate, and acetic acid, all LC-MS grade, were purchased from Fisher Scientific (Pittsburgh, PA). Ammonium hydroxide was bought from Sigma-Aldrich (Saint Louis, MO). DI water was provided in-house by a Water Purification System from EMD Millipore (Billerica, MA). PBS was bought from GE Healthcare Life Sciences (Logan, UT). The standard compounds corresponding to the measured metabolites were purchased from Sigma-Aldrich (Saint Louis, MO) and Fisher Scientific (Pittsburgh, PA).

Zebrafish intestine tissue samples (∼10-12 mg per sample) were homogenized in 200 µL MeOH:PBS (4:1, v:v, containing 1,810.5 μM ^13^C_3_-lactate and 142 μM ^13^C_5_-glutamic Acid) in an Eppendorf tube using a Bullet Blender homogenizer (Next Advance, Averill Park, NY). Then 800 µL MeOH:PBS (4:1, v:v, containing 1,810.5 μM ^13^C_3_-lactate and 142 μM ^13^C_5_-glutamic Acid) was added, and after vortexing for 10 s, the samples were stored at - 20°C for 30 min. The samples were then sonicated in an ice bath for 30 min. The samples were centrifuged at 14,000 RPM for 10 min (4°C), and 800 µL supernatant was transferred to a new Eppendorf tube. The samples were then dried under vacuum using a CentriVap Concentrator (Labconco, Fort Scott, KS). Prior to MS analysis, the obtained residue was reconstituted in 150 μL 40% PBS/60% ACN. A quality control (QC) sample was pooled from all the study samples.

The untargeted LC-MS metabolomics method used here was modeled after protocols reported in a number of studies^118-124^. Briefly, all LC-MS experiments were performed on a Thermo Vanquish UPLC-Exploris 240 Orbitrap MS instrument (Waltham, MA). Each sample was injected twice, 10 µL for analysis using negative ionization mode and 4 µL for analysis using positive ionization mode. Both chromatographic separations were performed in hydrophilic interaction chromatography (HILIC) mode on a Waters XBridge BEH Amide column (150 x 2.1 mm, 2.5 µm particle size, Waters Corporation, Milford, MA). The flow rate was 0.3 mL/min, auto-sampler temperature was kept at 4°C, and the column compartment was set at 40 °C. The mobile phase was composed of Solvents A (10 mM ammonium acetate, 10 mM ammonium hydroxide in 95% H_2_O/5% ACN) and B (10 mM ammonium acetate, 10 mM ammonium hydroxide in 95% ACN/5% H_2_O). After the initial 1 min isocratic elution of 90% B, the percentage of Solvent B decreased to 40% at t=11 min. The composition of Solvent B maintained at 40% for 4 min (t=15 min), and then the percentage of B gradually went back to 90%, to prepare for the next injection. Using mass spectrometer equipped with an electrospray ionization (ESI) source, we will collect untargeted data from 70 to 1050 m/z.

To identify peaks from the MS spectra, we made extensive use of the in-house chemical standards (∼600 aqueous metabolites), and in addition, we searched the resulting MS spectra against the HMDB library, Lipidmap database, METLIN database, as well as commercial databases including mzCloud, Metabolika, and ChemSpider. The absolute intensity threshold for the MS data extraction was 1,000, and the mass accuracy limit was set to 5 ppm. Identifications and annotations used available data for retention time (RT), exact mass (MS), MS/MS fragmentation pattern, and isotopic pattern. We used the Thermo Compound Discoverer 3.3 software for aqueous metabolomics data processing. The untargeted data were processed by the software for peak picking, alignment, and normalization. To improve rigor, only the signals/peaks with CV < 20% across quality control (QC) pools, and the signals showing up in >80% of all the samples were included for further analysis.

### Metabolomics Data Analysis

Metabolomics data preprocessing and analysis were performed using the online platform MetaboAnalyst 6.0^49^ (https://www.metaboanalyst.ca). Pathway enrichment analysis was conducted within MetaboAnalyst using the Enrichment Analysis module to identify significantly metabolic pathways. For visualization, heatmaps of selected metabolites were generated in R studio using the ComplexHeatmap (v2.20.0) packages.

### Fecal Matter Collection

At 25 dpf, following the morning feeding, zebrafish were transferred from nursery tanks into petri dishes containing system water at a density of 50 fish per dish. After three hours, fish were removed from the petri dishes. Water containing the fecal matter was collected and centrifuged. The supernatant was discarded, and the fecal matter pellet was immediately flash-frozen in liquid nitrogen and stored at -80°C.

### Metagenomics

Microbial DNA was isolated from frozen zebrafish fecal matter using Quick-DNA Fecal/Soil Microbe Microprep Kit (Zymo Research, Irvine, California) by following the manufacturer’s protocol. DNA concentrations were quantified using a NanoDrop 1000 Spectrophotometer (Thermo Scientific, Waltham, Massachusetts) at 260 nm. Sample QC was performed with Microplate Reader for concentration and Agarose Gel Electrophoresis for integrity. DNA samples was fragmented by ultrasound using the Covaris instrument. Short DNA fragments meeting the target length requirements (300-400 base pairs) were obtained by adjusting the breaking parameters. The fragmented samples were selected using the Agencourt AMPure XP-Medium kit, and the sample bands were concentrated at ∼ 300-400 bp. Purified DNA samples were quantified using the Qubit dsDNA HS Assay Kit 500 assays kit. The double-stranded DNA ends were repaired, and the “A” base was added at the 3’ end. An adapter was connected to DNA. Ligation products were amplified by PCR. The amplified products were subjected to fragment screening using Agencourt AMPure XPMedium. The PCR products were detected with an Agilent 2100 Bioanalyzer. After the PCR products are denatured into single stranded DNA, a cyclization reaction was performed to obtain a single-chain circular product. A final library is obtained after the uncyclized linear DNA molecules were removed by enzyme digestion. Fragment size and concentration of the library were detected using an Agilent 2100 bioanalyzer (Agilent DNA 1000 Reagents). Single-stranded circular DNA molecules were replicated by rolling circle amplification to obtain DNBs that each contained more than 300 copies of the molecule. The DNBs were loaded into a patterned nanoarray to acquire 150 bp paired-end reads. The sequencing was conducted on the MGI T7 platform.

### Metagenomics Data Analysis

We obtained 150 bp paired-end sequencing data for each sample. Initial quality control was performed using FastQC to assess read quality, and all samples were deemed to be of high quality. To remove host-derived sequences, raw FASTQ files were aligned to the *Danio rerio* reference genome (GRCz11) using Bowtie 2^125^ with default parameters. Only unmapped reads—those that did not align to the zebrafish genome—were retained for downstream microbial taxonomic analysis. On average, we retained approximately 20 million non-zebrafish reads per sample. We inferred the microbial origin of the remaining reads using Kraken 2^126,127^ (v2.1.3) with the Standard Kraken database. Kraken was run with default settings except for a modified classification threshold requiring a minimum of three overlapping k-mers to assign a read to a taxon, which is slightly more conservative than the default setting (two k-mers). This modification was made to increase classification specificity while maintaining sensitivity. Across samples, at least 40% of the non-host reads were classified as bacterial. To estimate read counts at different taxonomic levels, we used Bracken^128^ (Bayesian Reestimation of Abundance after Classification with Kraken) to re-estimate species, genus, and higher-level abundances from Kraken output files.

For differential abundance and prevalence analyses between treatment groups (CPF vs. DMSO), we used the MaAsLin 3 (Microbiome Multivariable Associations with Linear Models) R package^129^, designed to model both abundance (based on read counts) and prevalence (based on presence/absence) while accounting for the compositional nature of microbiome data. Prior to modeling, abundance data were normalized using total sum scaling (TSS), dividing each feature count by the total count per sample to account for differences in library size. The normalized dataset was split into two matrices: a binary prevalence matrix (presence/absence) and a non-zero abundance matrix for quantitative analysis. All metagenomics data analyses were conducted using R version 4.4.0 (2024-04-24) on a CentOS Linux 7 (Core) platform (x86_64-pc-linux-gnu).

### Griess Assay

To measure nitrite levels following CPF exposure, 85 embryos were exposed to 15 µM CPF or DMSO control from 0 to 3 dpf and collected at 6 dpf. After being weighed, larvae were washed, homogenized in 100 µL water, and centrifuged to collect the supernatants. The Griess assay was performed using the Griess Reagent Nitrite Measurement Kit (Cell Signaling Technology, #13547) according to the manufacturer’s instructions. Each group was analyzed in triplicate.

## ACKNOWLEDGEMENTS

We thank the University of Washington Office of Comparative Medicine for providing zebrafish husbandry support. This work was supported by the National Institute of Environmental Health Sciences (NIEHS) of the NIH under the award numbers R00ES031050, R01ES030197, R01ES031098, R25ES025503, and P30ES007033, and the University of Washington Environmental Health and Microbiome Research Center (EHMBRACE). The content is solely the responsibility of the authors and does not necessarily represent the official views of the NIH.

## AUTHOR CONTRIBUTIONS

L.D., A.X.K., and P.Z. designed and conducted the experiments and analyzed data. J.C. and H.G. conducted metabolomics analysis. K.P. M.J., A.E., I.D., and Y.M. assisted in conducting experiments and data analysis. J.W.M and T.B. analyzed metagenomics data. J.Y.C., H.G., T.B., and Y.G. planned the multi-omics analyses. Y.G. conceived the study, interpreted the data, and wrote the manuscript.

## DATA AND MATERIALS AVAILABILITY

All data needed to evaluate the conclusions in the paper are present in the paper and/or the Supplementary Materials.

