## Supplementary Materials for "Butyrate rescues chlorpyrifos-induced social deficits through inhibition of class I histone deacetylases"

**This PDF file includes:**

Supplementary Figures 1 to 14

Supplementary Tables 1 to 3 Legends

#### SUPPLEMENTARY FIGURES

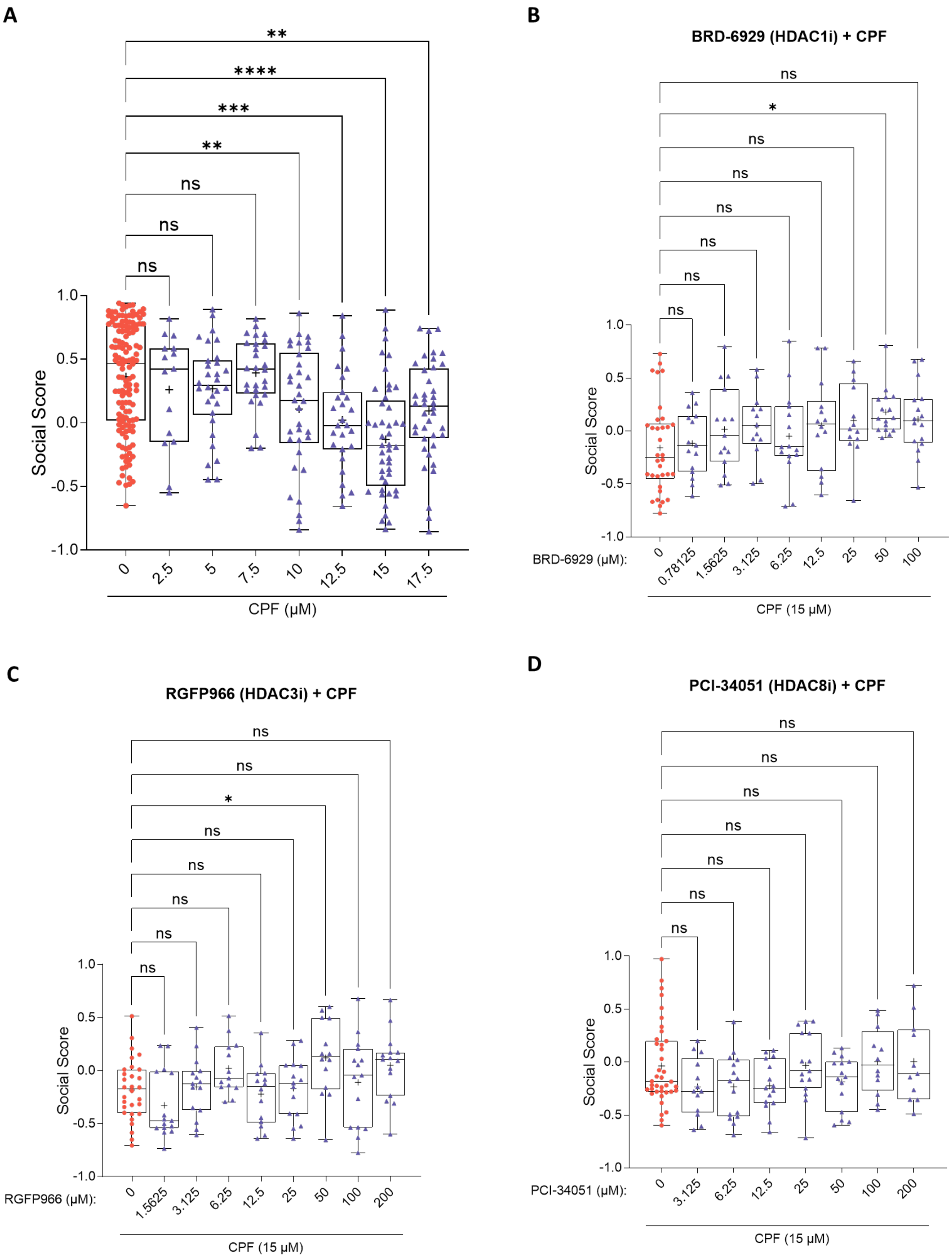


**Supplementary Figure 1.**

(A) Boxplot showing individual fish's social scores for the CPF dose curve. Each fish's social score is represented by a colored dot.

(B-D) Boxplot showing individual fish's social scores for the BRD-6929 (B), RGFP966 (C), and PCI-34051 (D) rescue experiments. All fish were pre-exposed to 15 μM CPF at 0-3 dpf.

Significance was calculated by one-way ANOVA and Dunnett's multiple comparison test. ns: not significant, **p*<0.05, ***p*<0.01, ****p*<0.001, *****p*<0.0001.

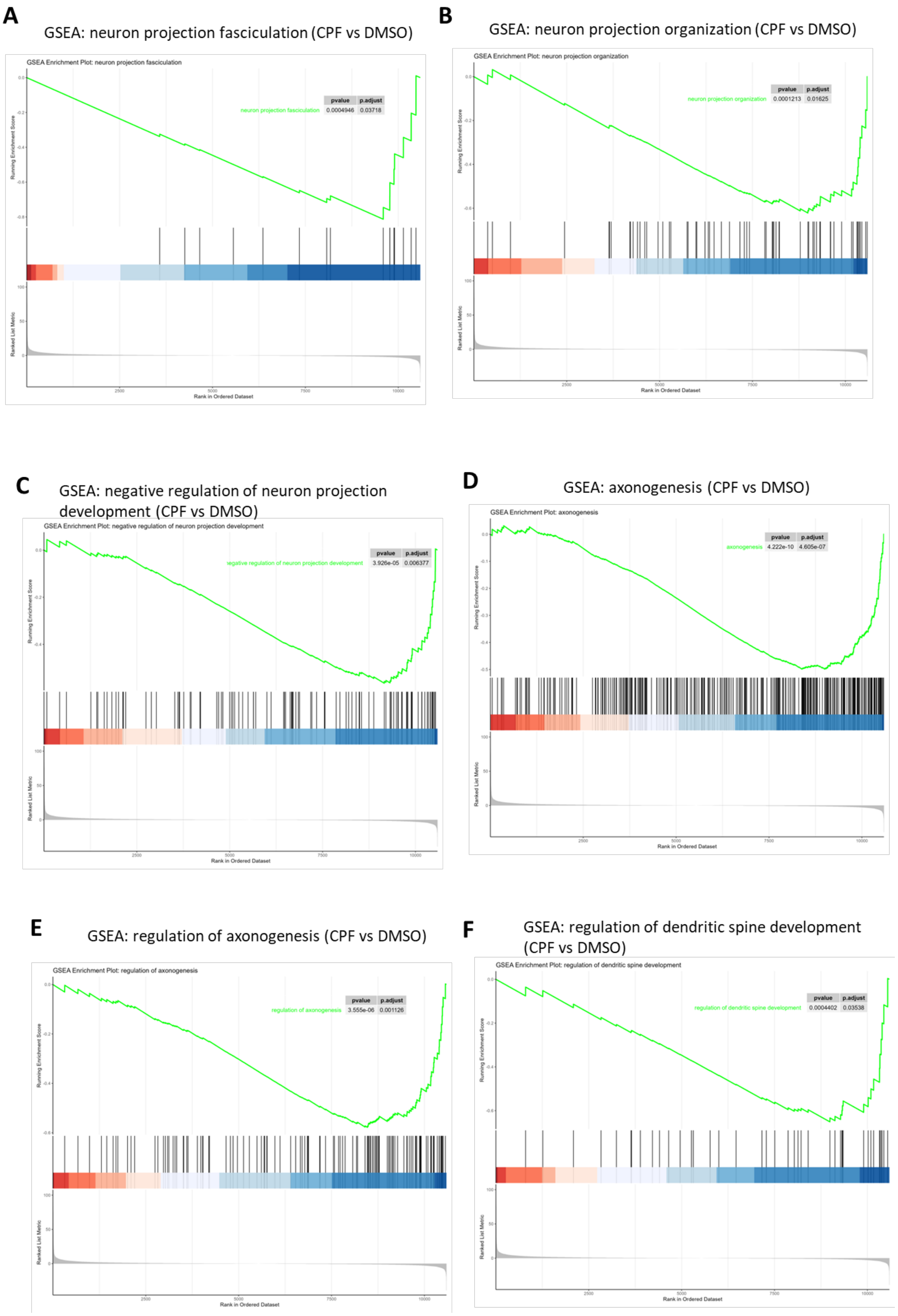


**Supplementary Figure 2.**

GSEA plots of pathways related to neuronal projection (A-C), axonogenesis (D-E), and dendrite development (F). Analyses were based on differential gene expression (RNA-seq results) between the CPF samples and the DMSO control samples.

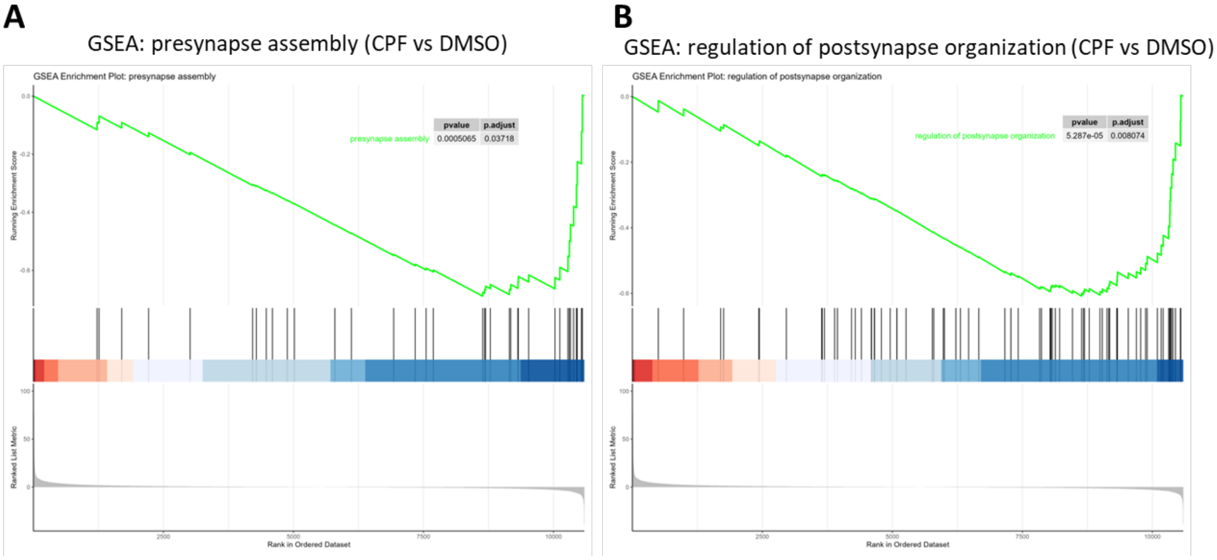


**Supplementary Figure 3.**

GSEA plots of pathways related to presynaptic (A) and postsynaptic (B) regulations. Analyses were based on differential gene expressions (RNA-seq results) between the CPF samples and the DMSO samples.

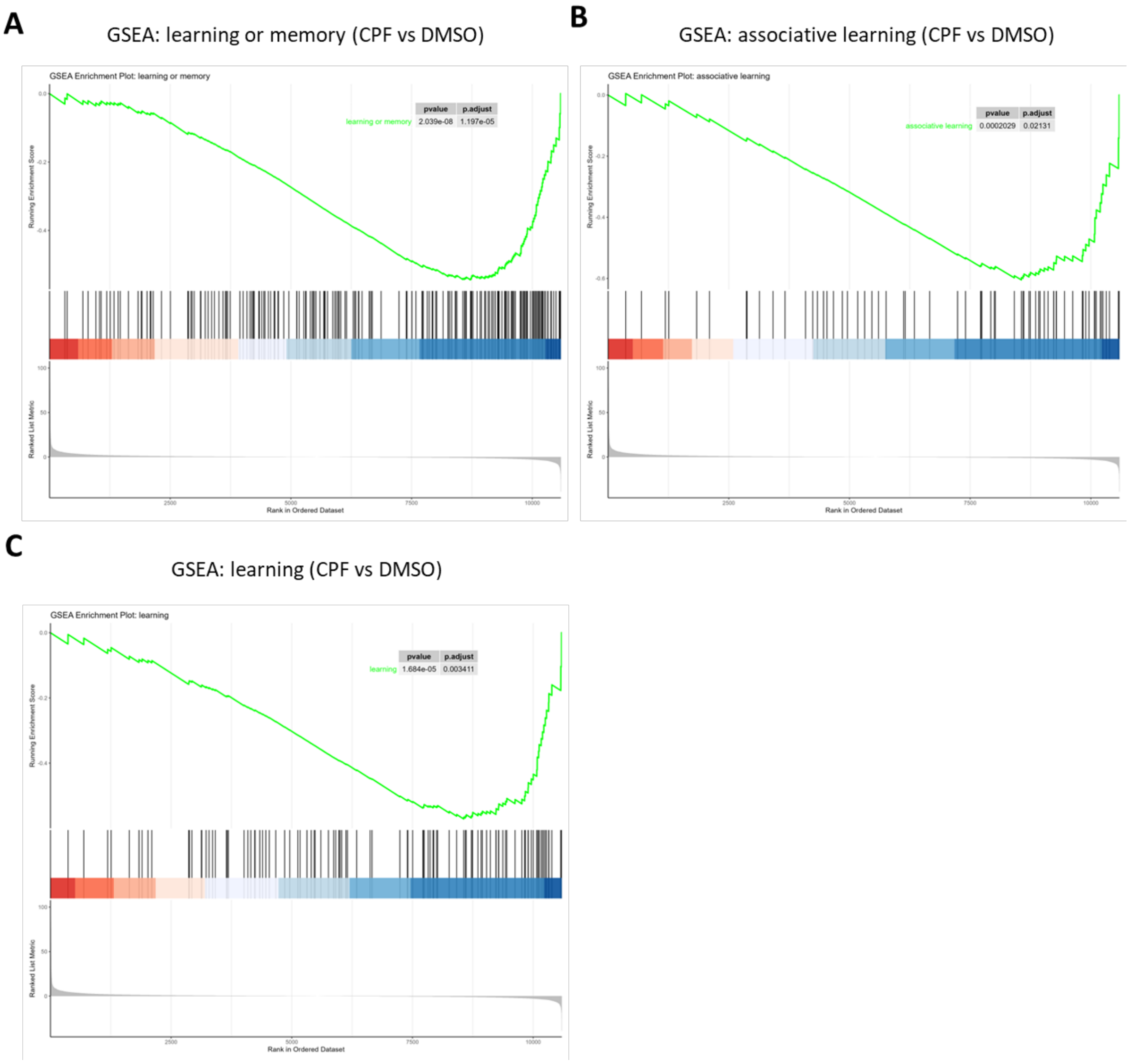


**Supplementary Figure 4.**

GSEA plots of pathways associated with learning (A-C). Analyses were based on differential gene expressions (RNA-seq results) between the CPF samples and the DMSO samples.

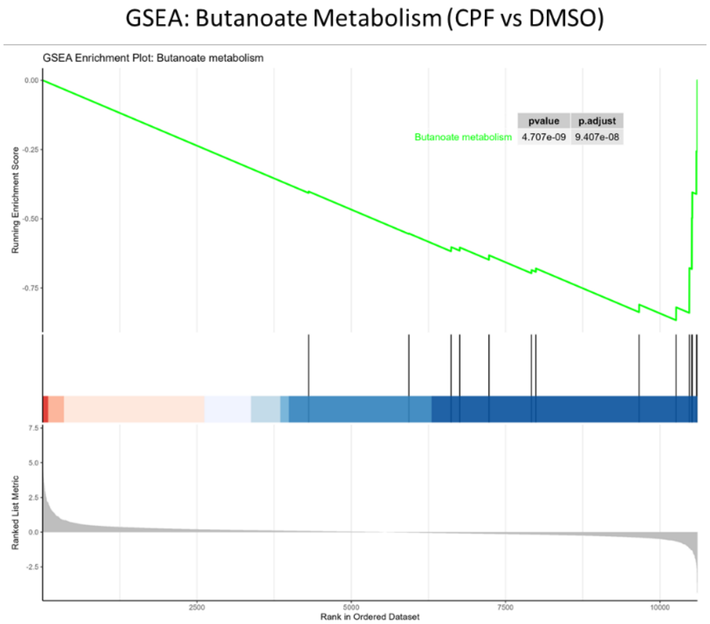


**Supplementary Figure 5.**

GSEA plot showing significant downregulation of the butanoate (butyrate) metabolism pathway in CPF samples as compared to DMSO control. Analysis was based on differential gene expression (RNA-seq results) between the CPF samples and the DMSO samples.

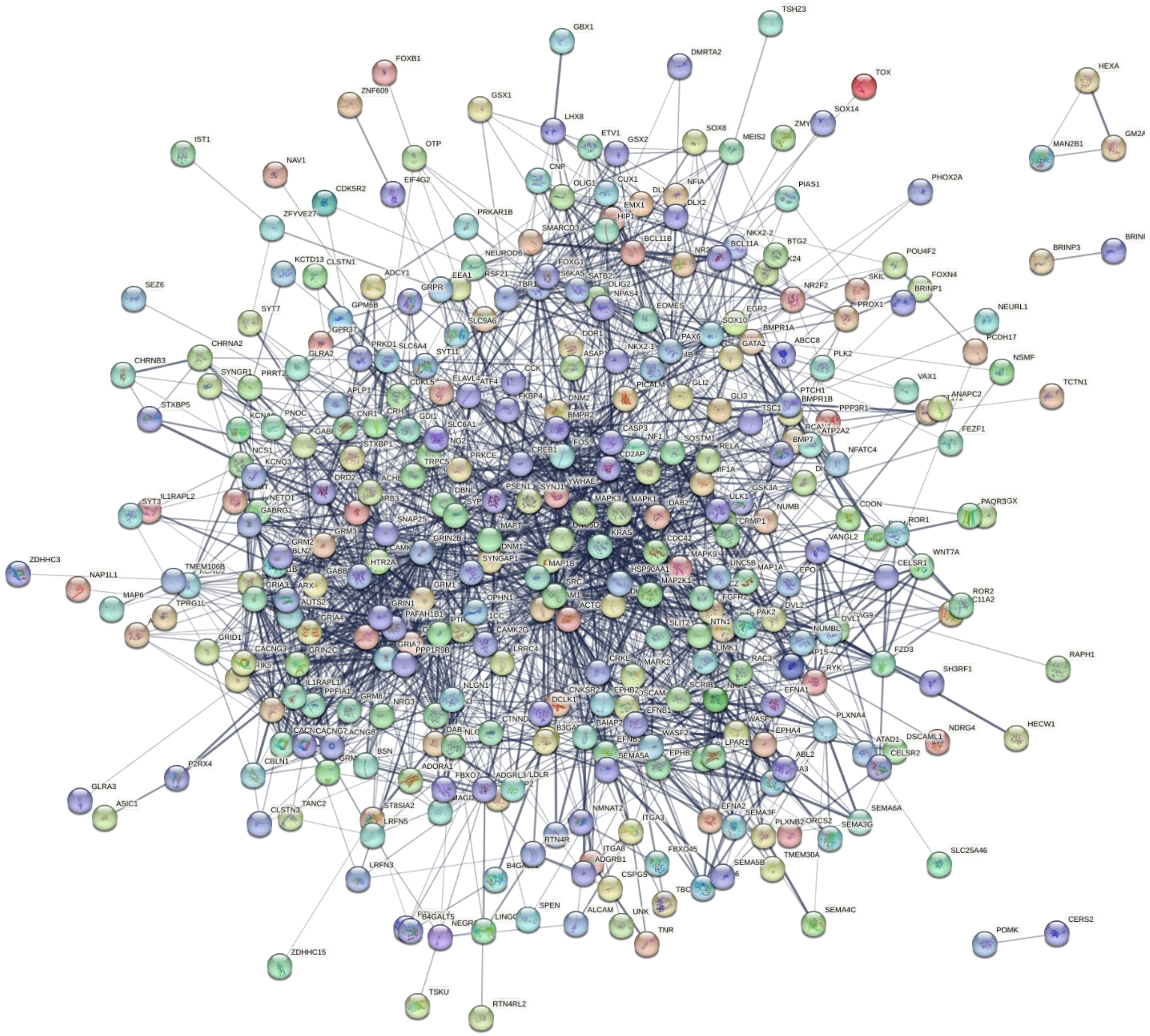


**Supplementary Figure 6.**

Protein-protein interaction (PPI) network formed by protein products of significant downregulated neuronal genes found in the CPF samples compared to the control samples. All genes belong to the 13 neuronal pathways found in Figure 4A.

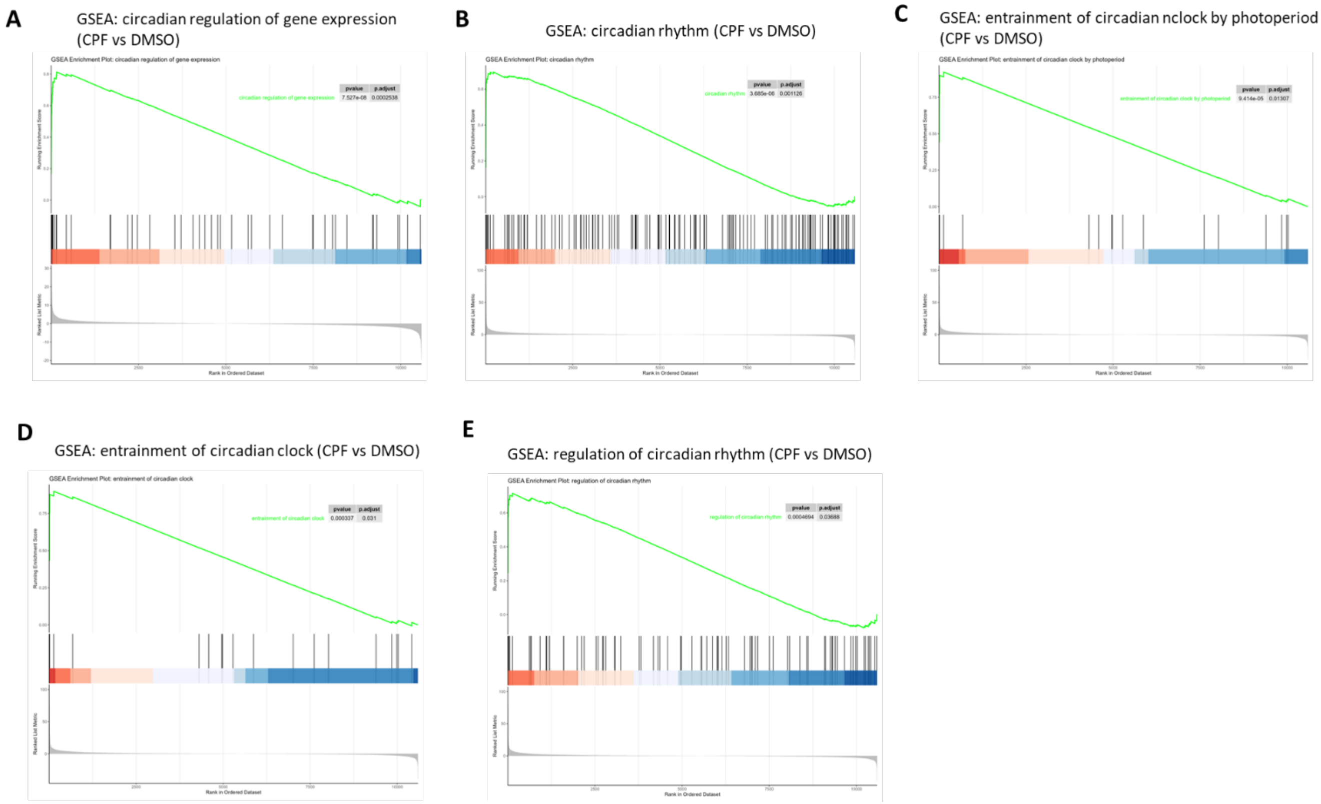


**Supplementary Figure 7.**

GSEA plot showing significant upregulation of circadian pathways (A-E) in CPF samples as compared to the DMSO control. Analyses were based on differential gene expressions (RNA-seq results) between the CPF samples and the DMSO samples.

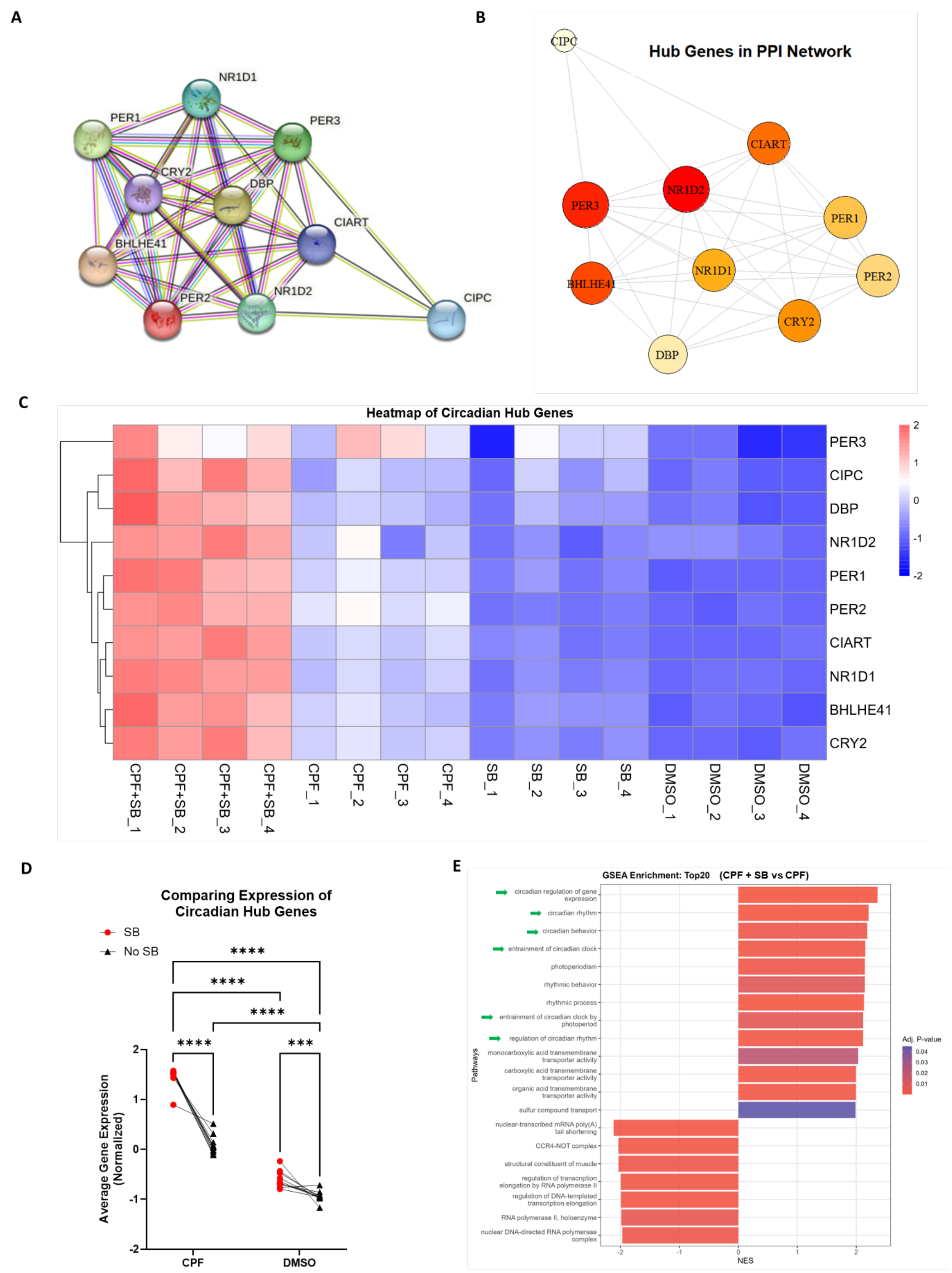


**Supplementary Figure 8.**

(A) Protein-protein interaction (PPI) network formed by protein products of significantly upregulated circadian genes found in the CPF samples compared to the control samples and belong to the 5 circadian pathways discovered by GSEA as shown in Figure 5A.

(B) Hub genes were identified by network topology analysis of the PPI network in (A). Genes were ranked by node degree values, with colors representing the rankings. Red denotes the top-ranking genes. The size of each colored circle represents the degree centrality (number of connections in the network).

(C) Gene expression heatmap shows upregulation of key circadian genes by CPF as compared to DMSO control (CPF vs. DMSO), and further upregulation by butyrate as compared to the CPF samples (CPF+SB vs. CPF).

(D) Comparing the average normalized expressions of circadian hub genes. Circadian hub gene expressions are significantly elevated in CPF samples compared to DMSO samples. Butyrate (SB) further upregulated the expression of circadian hub genes in CPF-treated fish. Lines between treatment groups connect expression levels of the same gene. Significance was calculated by two-way ANOVA with Tukey's multiple comparison test. ***$p < 0.001$, ****$p < 0.0001$.

(E) GSEA analysis comparing the gene expression profiles (RNA-seq results) of CPF+SB samples and CPF samples using Gene Ontology's (GO) Biological Process (BP), Molecular Function (MF), and Cellular Component (CC) terms. The top 20 pathways ranked by absolute values in Normalized Enrichment Score (NES) are shown, including 13 upregulated pathways and 7 downregulated pathways. 6 out of the 13 upregulated pathways are associated with circadian regulation (green arrows).

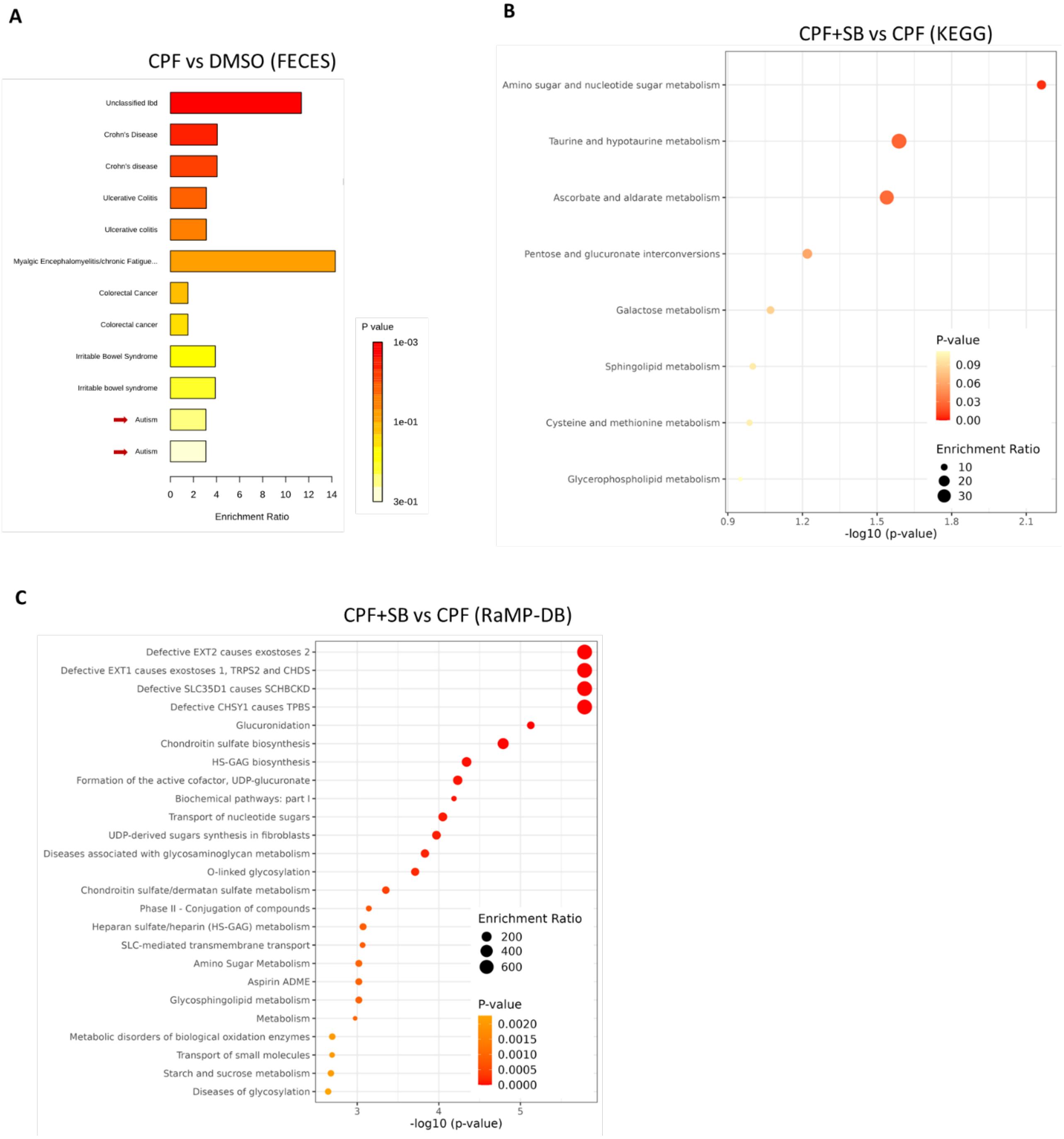


**Supplementary Figure 9.**

(A) Enrichment analysis comparing metabolomic changes in CPF samples as compared to DMSO controls, based on untargeted metabolomics analysis of gastrointestinal tissues and gut contents. Surveying the Feces dataset in HMDB detected similar patterns of metabolite profile changes in CPF-treated samples compared to fecal samples collected from individuals with inflammatory bowel disease (IBD) including Crohn's disease and colitis, irritable bowel syndrome (IBS), and autism (red arrows).

(B-C) Enrichment analysis comparing metabolomic changes between CPF and DMSO samples based on untargeted metabolomics analysis of gastrointestinal tissues and gut contents. Pathways related to the urea cycle, arginine metabolism, and aspartate metabolism were not identified by surveying KEGG (B) or RaMP-DB (C).

A

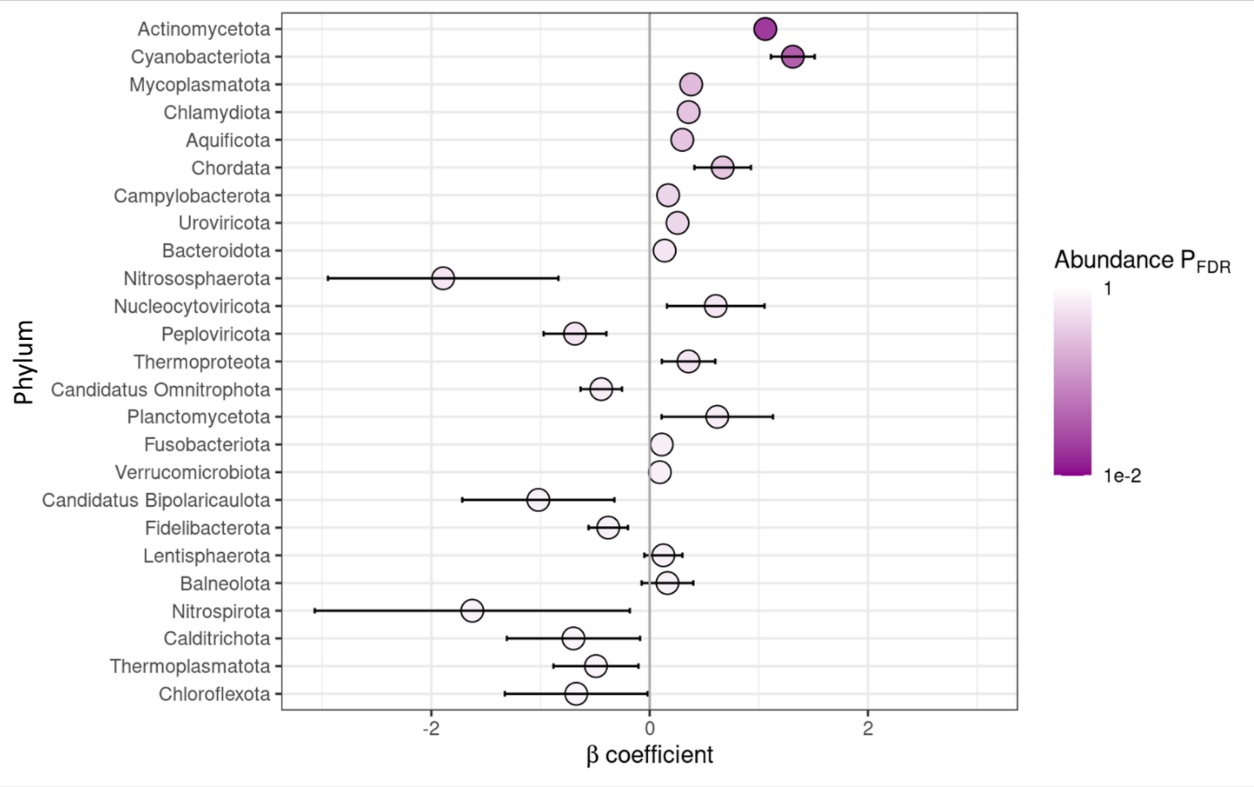


B

| Phylum | Coefficient | Std.Err | FDR |
|---|---:|---:|---:|
| *Actinomycetota* | 1.06 | 0.07 | 0.020 |
| *Cyanobacteriota* | 1.31 | 0.20 | 0.039 |

**Supplementary Figure 10.**

(A) β coefficient estimates and significance for the top 25 phyla. The plot presents the β coefficient for the test of CPF vs DMSO (the beta value is the log fold change), as well as the confidence interval for the coefficient (error bars). Colors of the symbols represent the FDR.

(B) Statistics for the two significant phyla.

A

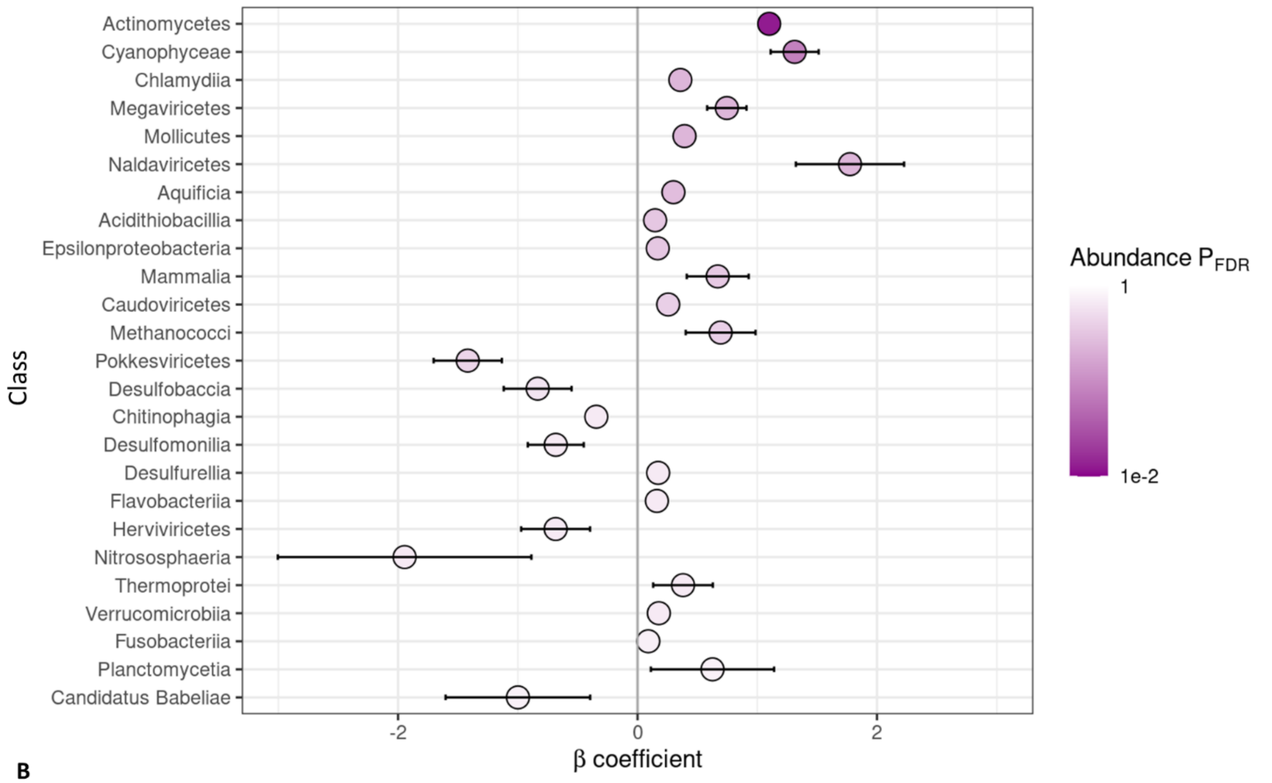


B

| Class | Coefficient | Std.Err | FDR |
|---|---:|---:|---:|
| *Actinomycetes* | 1.10 | 0.07 | 0.012 |
| *Cyanophyceae* | 1.31 | 0.20 | 0.078 |

**Supplementary Figure 11.**

(A) β coefficient estimates and significance for the top 25 classes. The plot presents the β coefficient for the test of CPF vs DMSO, as well as the confidence interval for the coefficient (error bars). Colors represent the FDR.

(B) Statistics for the two significant classes.

**A**

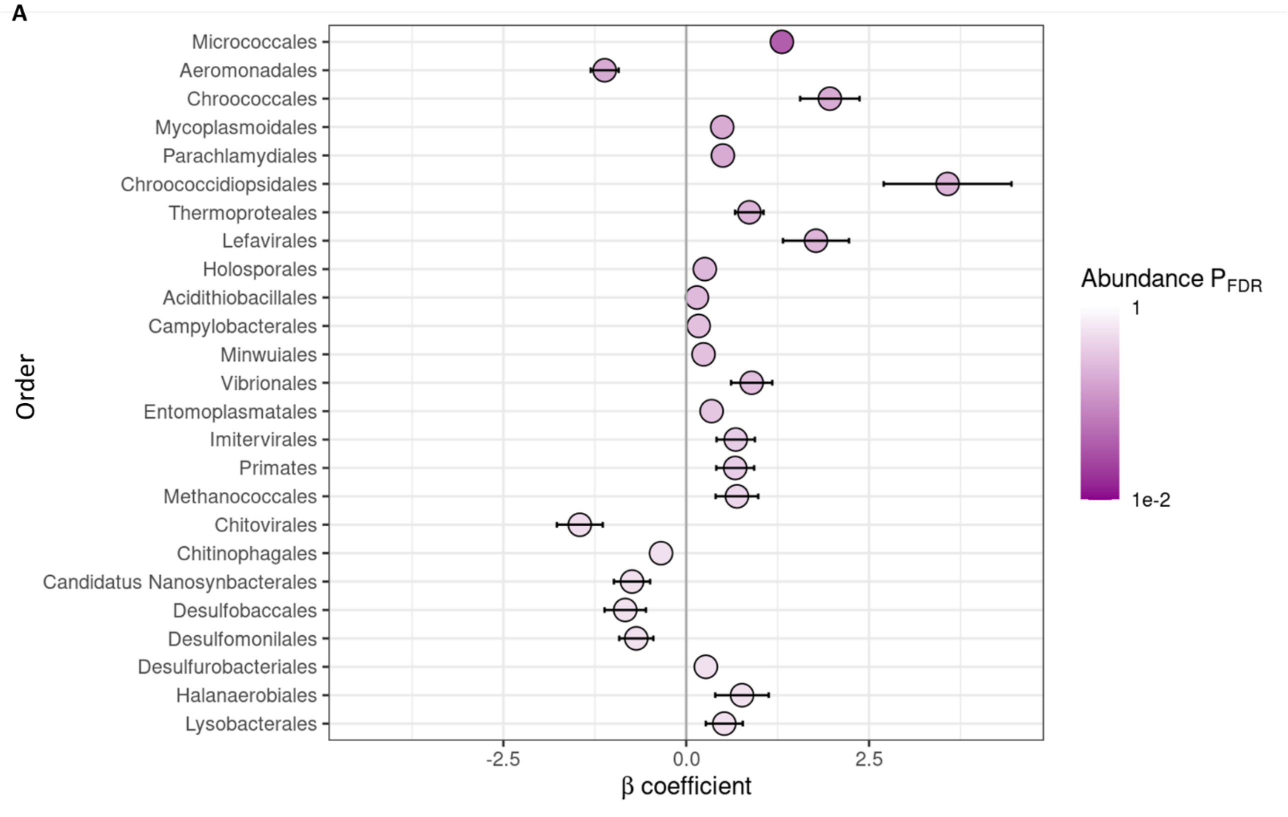


**B**

| Order | Coefficient | Std.Err | FDR |
|---|---:|---:|---:|
| *Micrococcales* | 1.31 | 0.11 | 0.039 |

**Supplementary Figure 12.**

(A) β coefficient estimates and significance for the top 25 orders. The plot presents the β coefficient for the test of CPF vs DMSO, as well as the confidence interval for the coefficient (error bars). Colors represent the FDR.

(B) Statistics for the significant order.

**A**

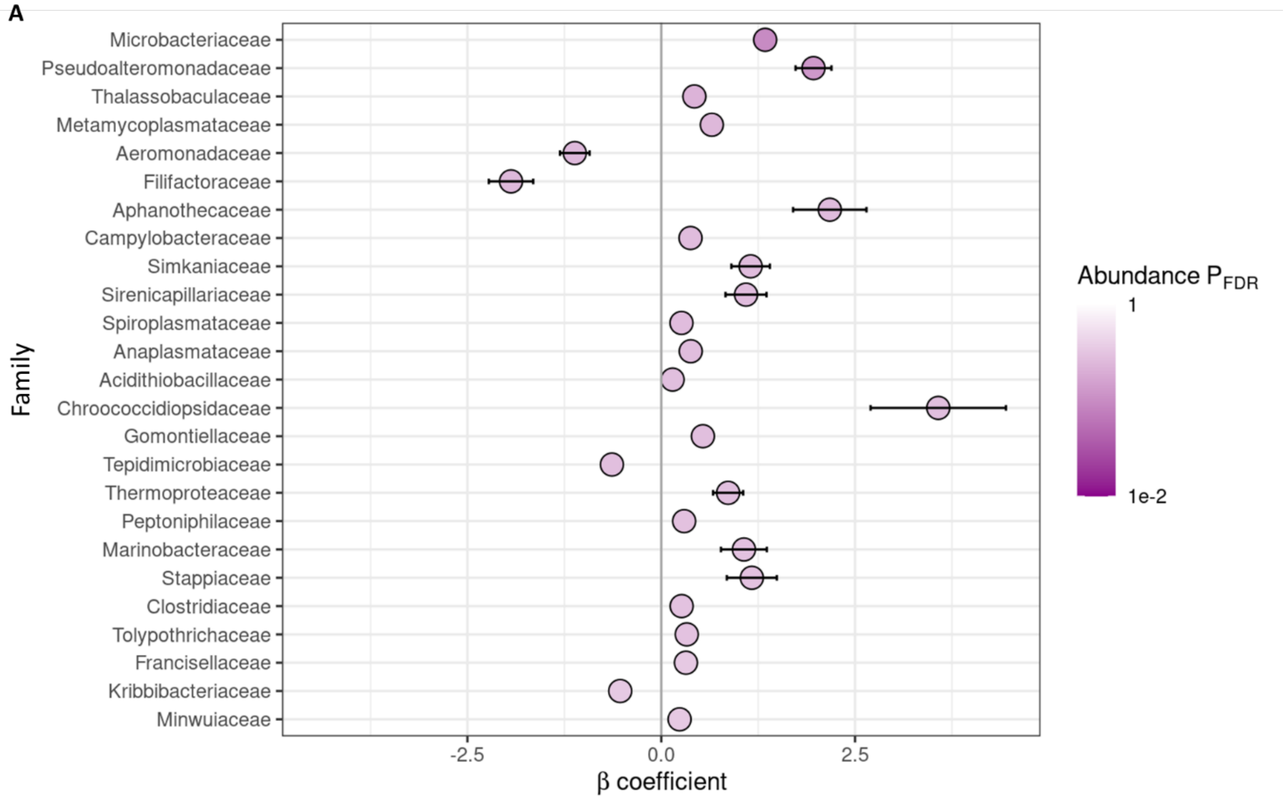


**B**

| **Family** | **Coefficient** | **Std.Err** | **FDR** |
|---|---:|---:|---:|
| *Microbacteriaceae* | 1.34 | 0.12 | 0.094 |

**Supplementary Figure 13.**

(A) β coefficient estimates and significance for the top 25 families. The plot presents the β coefficient for the test of CPF vs DMSO, as well as the confidence interval for the coefficient (error bars). Colors represent the FDR.

(B) Statistics for the significant family.

**A**

| Genus | Count |
|---|---|
| *Microbacterium* | 86 |
| *Aeromonas* | 11 |
| *Pseudomonas* | 6 |
| *Shewanella* | 3 |
| *Streptomyces* | 3 |
| *Labrenzia* | 2 |
| *Pseudoalteromonas* | 2 |
| *Vibrio* | 2 |
| *Alteromonas* | 1 |
| *Brenneria* | 1 |
| *Candidatus* | 1 |
| *Castellaniella* | 1 |
| *Corynebacterium* | 1 |
| *Ectopseudomonas* | 1 |
| *Edwardsiella* | 1 |
| *Geminocystis* | 1 |
| *Halomonas* | 1 |
| *Kocuria* | 1 |
| *Lacinutrix* | 1 |
| *Leifsonia* | 1 |
| *Leptospira* | 1 |
| *Nonlabens* | 1 |
| *Polaribacter* | 1 |
| *Pseudochrobactrum* | 1 |
| *Pseudoduganella* | 1 |
| *Pseudovibrio* | 1 |
| *Rathayibacter* | 1 |
| *Subtercola* | 1 |
| *Synechococcus* | 1 |
| *Thalassobaculum* | 1 |
| *uncultured* | 1 |
| *Xanthomonas* | 1 |

**B**

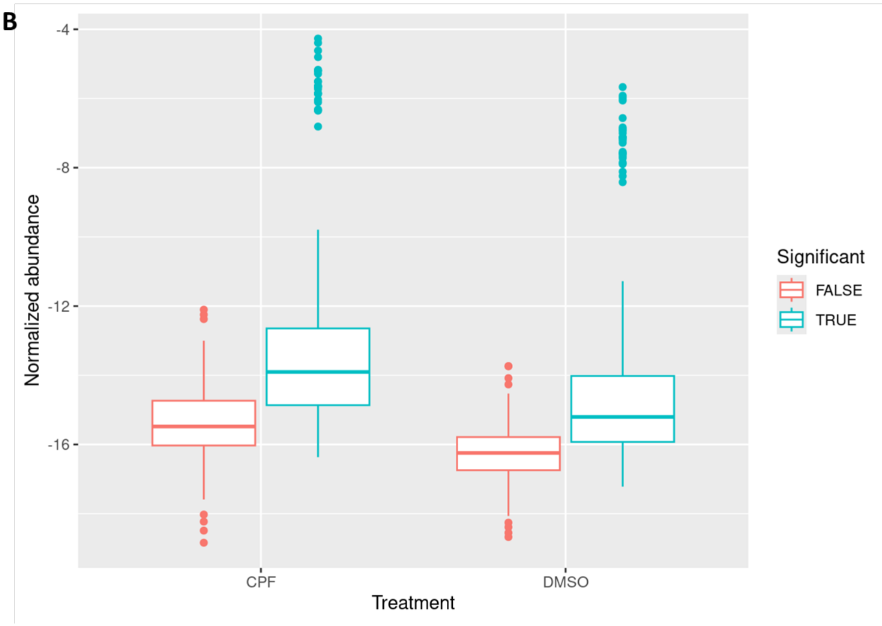


**Supplementary Figure 14.**

(A) Counts of all differentially abundant species by genus.

(B) Boxplots of the normalized abundance measures for all detected *Microbacterium* species. Data points were grouped by significance. The abundances of the non-significant species are similar between CPF and DMSO, but much lower compared to that of the significant species.

### SUPPLEMENTARY TABLE LEGENDS

**Supplementary Table 1.**

β coefficient estimates for 139 significant species detected by comparing microbiome compositions of fecal matters collected from CPF-treated fish and DMSO control fish. The microbiome profile of each sample was identified by shotgun metagenomics analysis.

**Supplementary Table 2.**

Design of gRNA sequences for knocking out the zebrafish *hdac1*, *hdac3*, *hdac4*, *hdac5*, *hdac8*, and *hdac9* genes.

**Supplementary Table 3.**

Design of quantitative PCR (qPCR) primer sequences for key zebrafish circadian rhythm genes.
